# Abnormal Lipid Metabolism and Altered Neuronal Support by Astrocytes Lacking *Akap11*, a Risk Gene for Schizophrenia and Bipolar Disorder

**DOI:** 10.1101/2025.04.25.650548

**Authors:** Xiao-Man Liu, Horia Pribiag, Deeksha Misri, Kwanho Kim, Kira Perzel Mandell, Bryan J. Song, David Graykowski, Olivia Seidel, Nolan D. Hartley, Mateo Herrera, He J. Xu, Matthew Tegtmeyer, Yan-Ling Zhang, Lei Cui, Natalie Clark, Wei-Chao Huang, John Adeleye, Courtney Dennis, Lucas Dailey, Amy Deik, Lucia Inunciaga, Eivgeni Mashin, Sean K Simmons, Jen Q. Pan, Ralda Nehme, Hasmik Keshishian, Steven A. Carr, Zhanyan Fu, Joshua Z. Levin, Clary B. Clish, Morgan Sheng

## Abstract

*A-Kinase Anchoring Protein 11* (*AKAP11*) is a shared genetic risk factor for schizophrenia and bipolar disorder, yet its role in the brain remains poorly understood. Through multi-omic analysis of *Akap11* mutant mouse brains and cultured astrocytes, we identified significant transcriptomic, proteomic, and metabolomic alterations. Key findings include the upregulation of cholesterol and fatty acid metabolic pathways, accumulation of lipid species such as cholesteryl esters, triacylglycerols, ceramides, and glycerophospholipids, and elevated levels of 3’,5’-cyclic AMP and protein kinase A (PKA) signaling. These metabolic perturbations manifested as increased lipid droplet accumulation in *Akap11* mutant astrocytes, highlighting AKAP11’s critical role in regulating intracellular lipid homeostasis. Mechanistically, AKAP11 functions as an autophagy receptor mediating PKA degradation and interacts with endoplasmic reticulum-resident proteins VAP-A and VAP-B through its FFAT motif, providing possible molecular insight into AKAP11’s regulation of lipid metabolism. Co-culture experiments with mouse astrocytes and human induced pluripotent stem cell-derived neurons demonstrated that loss of *Akap11* in astrocytes, relative to wild-type, increases excitatory neurotransmission and neuronal activity. Collectively, these findings link AKAP11-mediated lipid and synaptic dysregulation to psychiatric disease risk and highlight the potential role of astrocytes in these disorders.

## Introduction

Schizophrenia (SCZ) and bipolar disorder (BD) are highly heritable psychiatric diseases that typically present in adolescents and young adults^1,2^. These disorders impose a heavy burden on affected individuals, their families, and society, with patients facing an average reduction in life expectancy of 10-20 years compared to the general population^3^. While the disease mechanisms of SCZ and BD remain largely unclear, the two disorders share mood and psychotic symptoms, and significant overlap in genetic risk factors^4–10^. Recent exome sequencing studies have uncovered a number of genes with rare coding variants that significantly increase the risk of developing SCZ or BD^11,12^. These rare variants, exhibiting large effect sizes (odds ratios ranging from 3 to 50) offer potential inroads into the disease mechanisms^13^. Among the identified genes, *AKAP11* (*A-Kinase Anchoring Protein 11*) has emerged as a rare variant risk gene for both SCZ and BD^12,14^.

AKAPs bind directly to the regulatory subunits of protein kinase A (PKA), anchoring the PKA holoenzyme to specific locations within the cell^15^. Besides PKA, AKAPs often interact with signal termination enzymes, such as phosphodiesterases (PDE) and phosphatases, to form AKAP signalosomes that help to establish subcellular compartments for cyclic adenosine 3′,5′-monophosphate (cAMP) signaling^16,17^. AKAP11 (also known as AKAP220) functions as a scaffolding protein that binds both PKA subunits and protein phosphatase 1 (PP1)^18,19^. Within cells, AKAP11 forms punctate structures and has been reported to localize to the endoplasmic reticulum (ER)^20^, peroxisomes^15^, multivesicular bodies^19^, and autophagosome^21^. While its role in these organelles remains largely unknown, AKAP11 has been shown to act as a selective autophagy receptor, facilitating the recruitment of the PKA subunit PRKAR1 to autophagosomes for degradation^21,22^. Beyond its organelle-associated functions, AKAP11 regulates diverse cellular processes through specific protein interactions: it controls microtubule dynamics via IQGAP1^23^, maintains renal water homeostasis through aquaporin-2 (AQP2)^24^, modulates endothelial barrier function through VE-cadherin/β-catenin^25^, and controls primary cilia development by associating with PP1^26^. AKAP11 also interacts with VAMP-associated proteins (VAPs) VAPA and VAPB^27^, although the biological significance of this interaction remains unclear.

In the context of psychiatric disorders, a particularly intriguing aspect of AKAP11 is its ability to interact with and inhibit glycogen synthase kinase 3 beta (GSK3β)^28^, as GSK3β is a major target of lithium, a mainstay treatment for BD^29^. We have previously reported that *Akap11* mutant mice display electroencephalogram (EEG) abnormalities that overlap those observed in SCZ patients^30^. To explore AKAP11’s function in the brain, we recently conducted detailed transcriptomic and proteomics analyses of *Akap11* mutant mouse brains^27^, showing widespread changes in gene expression in neurons and glia, and markedly elevated PKA protein levels and PKA activity at synapses. *Akap11* mRNA is expressed in neurons as well as astrocytes across the brain. In the prefrontal cortex (PFC) at four weeks, *Akap11*^+/-^ astrocytes exhibited a particularly high number of differentially expressed genes (DEGs)^27^, which hints at a pivotal role for AKAP11 in astrocytic function. Moreover, *Akap11* loss of function in astrocytes resulted in significant downregulation of many genes involved in the sterol/cholesterol biosynthetic pathway^27^.

In this study, we sought to elucidate the role of AKAP11 in astrocytes and its broader implications for brain function. By employing an integrative multi-omic approach, we examined transcriptomic, proteomic, and metabolomic alterations caused by *Akap11* loss-of-function in astrocytes. *Akap11*-deficient astrocytes show significant upregulation of lipid metabolism, along with elevated levels of 3’,5’-cyclic AMP and PKA subunits. We further investigated the molecular mechanisms underlying these alterations by examining AKAP11’s role in autophagic degradation of PKA and its interaction with ER-associated VAP proteins. Furthermore, we demonstrated that astrocytic AKAP11 dysfunction significantly increases neuronal activity in coculture experiments. Collectively, these results provide mechanistic insights into AKAP11’s function in astrocytes, revealing a pivotal role in regulating various aspects of cellular metabolism, and suggesting a potentially crucial role for astrocytic dysfunction in the pathophysiology of SCZ and BD.

## Results

### Transcriptomic and proteomic analysis reveals upregulated lipid pathways in *Akap11* mutant astrocytes

To elucidate AKAP11’s role in astrocyte function, we conducted transcriptomic and proteomic analyses on cultures of primary astrocytes isolated from the cerebral cortex of *Akap11*^+/+^, *Akap11*^+/-^, and *Akap11*^-/-^ mouse pups (Figure 1A). Differential expression analysis of bulk RNA-seq of cultured astrocytes revealed numerous DEGs (FDR < 0.05) in both *Akap11*^+/-^ and *Akap11*^-/-^ astrocytes (Figure S1A), with moderately correlated gene expression changes between the two genotypes (Pearson’s r = 0.48; Figure S1B).

**Figure 1.**
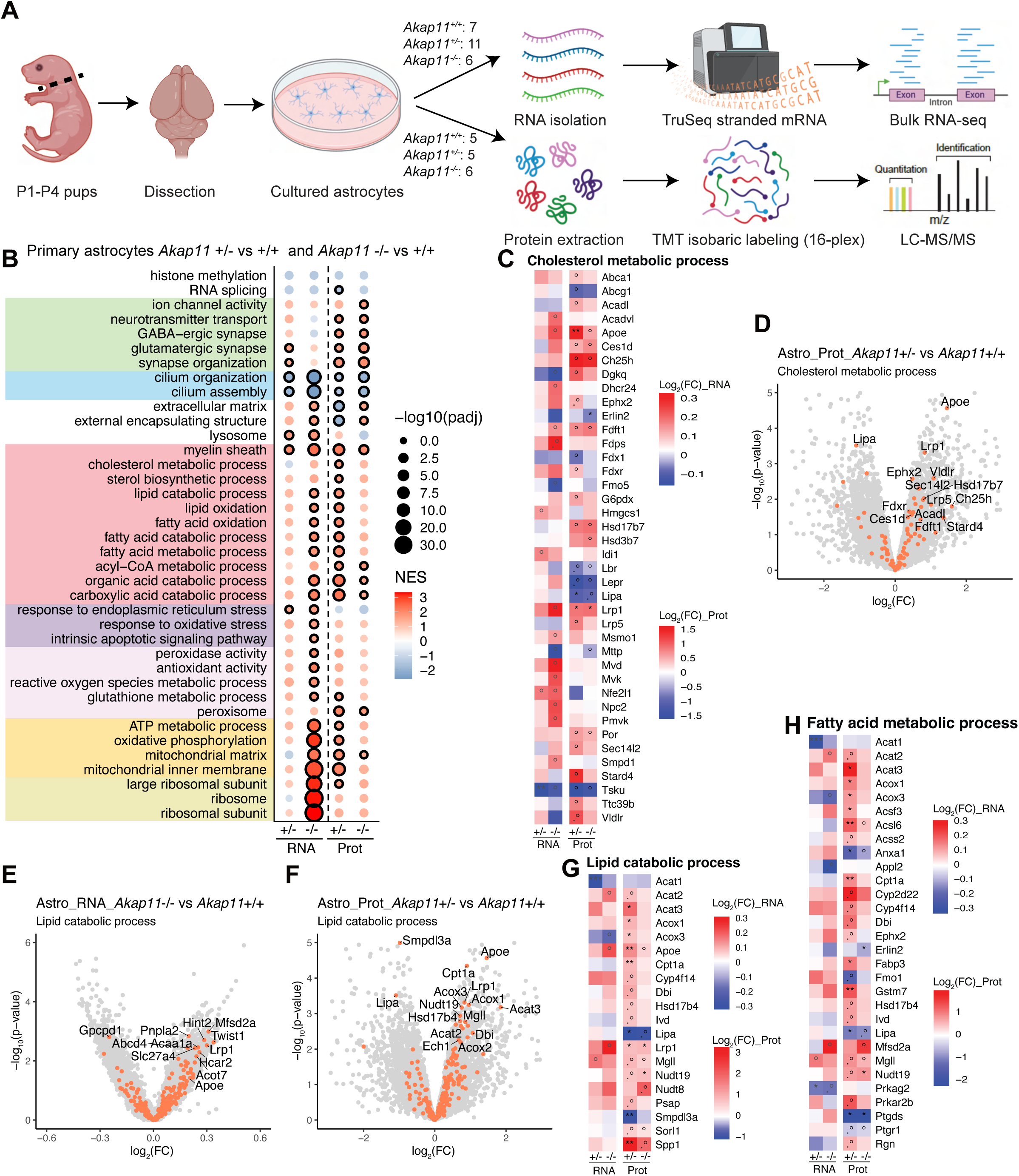
Transcriptomic and proteomic changes in multiple molecular pathways in *Akap11* mutant astrocytes. **(A)** Schematic overview of bulk RNA sequencing and proteomic analysis performed on primary astrocytes isolated from *Akap11*^+/+^, *Akap11*^+/-^, and *Akap11*^-/-^ mice at postnatal days 1-4 (P1-P4). Astrocytes were cultured for 12 days *in vitro*, replated, and cultured for an additional 3 days in serum-free media. RNA libraries were prepared with the Illumina TruSeq Stranded mRNA kit and sequenced on the NovaSeq S2 platform. For proteomic analysis, astrocytes were lysed, digested, labeled with tandem mass tag (TMT) reagents, and analyzed via liquid chromatography-tandem mass spectrometry (LC-MS/MS). All experiments and analyses were conducted in a blinded manner. The number of mice used for RNA sequencing and proteomics is indicated. **(B)** Gene set enrichment analysis (GSEA) reveals significant changes in multiple molecular pathways in *Akap11*^+/-^ and *Akap11*^-/-^ astrocytes compare to *Akap11*^+/+^. Bubble color represents the normalized enrichment score, with circles outlined in black indicating statistical significance (FDR < 0.05). **(C)** Heatmaps displaying log_2_FC values for genes and proteins involved in cholesterol metabolic process. Heatmaps include only those with a p-value < 0.05 in at least one condition. Significance levels are indicated as follows: padj ≤ 0.1, “.”; padj < 0.05, “*”; padj < 0.01, “**”; nominal p-value < 0.05: “°”. **(D-F)** Volcano plots highlighting transcriptomic and proteomic changes in *Akap11*^+/-^ and *Akap11*^-/-^ astrocytes compare to *Akap11*^+/+^, focusing on genes and proteins involved in cholesterol metabolic process (**D**) and lipid catabolic process (**E and F**). Orange dots indicate genes or proteins within the respective gene sets. **(G and H)** Heatmaps displaying log_2_FC values for genes and proteins involved in lipid catabolic process (**G**) and fatty acid metabolic process (**H**). Heatmaps include only those with a padj < 0.1 in at least one condition. Significance levels are indicated as follows: padj ≤ 0.1, “.”; padj < 0.05, “*”; padj < 0.01, “**”; padj < 0.001, “***”; nominal p-value < 0.05, “°”.

In parallel, we conducted mass spectrometry-based quantitative proteomics of astrocytes using multiplexed tandem mass tag (TMT16) labeling, which identified 133,247 unique peptides corresponding to 10,137 proteins. Differential expression analysis revealed numerous differentially expressed proteins (FDR < 0.05) in *Akap11*^+/-^ and *Akap11*^-/-^ astrocytes (Figure S1A) that were well correlated between genotypes (Pearson’s r = 0.62; Figure S1B).

To identify the molecular pathways affected by *Akap11* deficiency, we performed gene set enrichment analysis (GSEA) on the transcriptomic and proteomic datasets (Figure 1B). Pathway-level analysis revealed a good correlation in perturbed pathways (FDR < 0.05) between transcriptome and proteome data in both *Akap11*^+/-^ and *Akap11*^-/-^ astrocytes (Figure S1C). These “convergent pathways” include not only synapses, oxidative phosphorylation (OXPHOS), ribosomes, and fatty acid and cholesterol metabolic processes—pathways previously shown to be dysregulated in SCZ brain tissue^31–33^— but also pathways related to cilium organization and cilium assembly, which were among the most significantly downregulated (Figure 1B).

Among the pathways significantly enriched in the upregulated gene set, the most prominent changes were related to cellular stress, energy metabolism, and mitochondrial function. This was evidenced by positive normalized enrichment scores (NES) for mitochondrial matrix, inner mitochondrial membrane, OXPHOS, and ATP production. We also noted significant upregulation of lipid-related pathways, including fatty acid and cholesterol metabolism, lipid oxidation, and acyl-CoA metabolism in *Akap11* mutant astrocytes. Consistent with enhanced lipid catabolism, we found enrichment of both mitochondrial and peroxisomal pathways, the primary sites of cellular lipid oxidation (Figure 1B). These extensive changes lipid metabolic pathway were accompanied by cellular stress responses, evidenced by upregulation of gene sets such as “response to endoplasmic reticulum stress” (eg., *Ube2g2*, *Rcn3*, *Derl1*, *Fbxo6*, *Ddrgk1*, *Calr*, and *Nfe2l1*), “response to oxidative stress” (eg., *Sod3*, *Gpx7*, *Gpx1*, *Apex1*, *Zfp622*, and *mt-Nd3*), “reactive oxygen species metabolic process” (eg., *Gadd45a*, *Cycs*, *Sod3*, *Eif5a*, *Lcn2*, *Gpx1*, *Dcxr*, *Rhoa*, *Prdx5*, and *Prdx2*), and “intrinsic apoptotic signaling pathway” (g., *Cycs*, *Bcl2a1d*, *Hspb1*, *Bcl2a1b*, *Phlda3*, and *Ddit3*) (Figure 1B and Table S1).

Because transcriptomic evidence for dysregulation of cholesterol and fatty acid biosynthesis pathways in astrocytes has been observed in the *Grin2a* mutant SCZ mouse model^31^ and in human postmortem SCZ brain tissues^33^, we further examined alterations in these pathways in our *Akap11* mutant astrocytes. Notably, apolipoprotein E (*ApoE*), which transports cholesterol/lipids between cells, and its receptor, LDLR-related protein 1 (*Lrp1*), were found to be upregulated at the RNA level, with a significant increase detected at the protein level in both *Akap11*^+/-^ and *Akap11*^-/-^ astrocytes (Figures 1C, 1D, and S1G). Other ApoE receptors, *Lrp5* and *Vldlr* also showed elevated protein levels in *Akap11*^+/-^ astrocytes, although these increases did not reach statistical significance (Figures 1C and 1D). Individual genes in the cholesterol synthesis pathway did not exhibit statistically significant changes (FDR < 0.05) in RNA-seq or proteomics of astrocytes, though many showed an upward trend (nominal p < 0.05) in *Akap11* mutant astrocytes (Figures S1D-S1F). On the other hand, proteins involved in cholesterol ester synthesis, such as Acyl-coenzyme A acyltransferase (ACAT) enzymes encoded by *Acat2* and *Acat3*^34^, were significantly increased, specifically in *Akap11*^+/-^ astrocytes (Figures 1F and 1G). Ch25h, which encodes cholesterol 25-hydroxylase and converts cholesterol to 25-hydroxycholesterol (25-HC), was upregulated in protein level (Figure 1C). This increase in 25-HC could promote cholesterol esterification in astrocytes^35^. In contrast, *Lipa*, a lysosomal acid lipase crucial for hydrolyzing cholesterol esters to free cholesterol, was markedly downregulated at the protein level (Figures 1C and 1D). In aggregate, these changes would be predicted to increase total cholesterol esters levels.

The transcriptomic and proteomic alterations extended to fatty acid metabolism, where key enzymes involved in beta-oxidation and fatty acid synthesis were significantly dysregulated. In *Akap11*^+/-^ astrocytes, proteins such as Acox1 and Acox3, which initiate peroxisomal fatty acid beta-oxidation, and Cpt1, the rate-limiting enzyme for transporting long-chain fatty acids to the mitochondria for β-oxidation, were markedly upregulated (Figures 1H and S1I). Additionally, the *Eci1* gene, encoding a mitochondrial enzyme essential for beta-oxidation of unsaturated fatty acids, showed increased expression specifically in *Akap11*^-/-^ astrocytes (Figure S1H). Beyond beta-oxidation, genes involved in fatty acid synthesis and transport were also affected. For instance, in *Akap11*^+/-^ astrocytes, Acsf3, which converts malonic acid to malonyl-CoA in mitochondria; Acss2, which synthesizes acetyl-CoA from short-chain fatty acids in the cytosol; and Acsl6, which catalyzes the conversion of long-chain fatty acids to their active form acyl-CoA, were all upregulated in the proteomics data (Figures 1H and S1I). Similarly, *Fasn*, which catalyzes fatty acid synthesis, *Mfsd2a*, a transporter for omega-3 fatty acids^36^, and *Fabp3*, a fatty acid binding protein that facilitates fatty acid transfer, were elevated in *Akap11* mutant astrocytes (Figures 1H and S1H-S1I). In summary, *Akap11* mutant astrocytes display predominantly upregulated expression of genes and proteins involved in cholesterol transport and esterification, and fatty acid synthesis and β-oxidation pathways.

### Single-cell transcriptomics reveals astrocyte lipid dysregulation in *Akap11*-deficient mice

To determine whether the molecular changes observed in cultured *Akap11* mutant astrocytes are reflective of *in vivo* conditions, we conducted single-cell RNA sequencing (scRNA-seq) on astrocytes directly isolated from the cerebrum of 1-month-old *Akap11*^+/+^, *Akap11*^+/-^, and *Akap11*^-/-^ mice (Figure 2A). Following quality control, the RNA sequencing data were analyzed using the Seurat R package to identify cell clusters and perform dimensionality reduction (see methods; Figure S2B). Despite some residual contamination from other CNS cell types, astrocytes accounted for 50%-75% of the MACS sorted cells across the samples (Figure S2C). Cell-type identification was guided by MapMyCells, an automated approach to label cells by hierarchical mapping to a reference data^37^ (see Methods), and the expression of established cell type-specific marker genes (Figures S2A and S2D). After excluding contaminating cell types and low-quality cells (Figures, S2E-S2H), a total of 87,746 astrocytes remained, which were then re-clustered into three astrocyte subtypes: Astro-OLF (olfactory areas), Astro-TE (telencephalon), and Astro-NT (non-telencephalon), as described previously^38^. Among these, Astro-TE was the predominant subtype, representing over 70% of the cells (Figure S3A). Previously identified region-specific astrocyte genes including *Ppp1r3g*, *Agt*, and *Islr* were primarily used to confirm the annotation of Astro-TE, Astro-NT, and Astro-OLF (Figure S3B). Due to the exclusion of olfactory regions from the dissection protocol, the Astro-OLF subcluster (comprising 4001 cells) was omitted from subsequent analyses (Figures 2B and S3C-3E).

**Figure 2.**
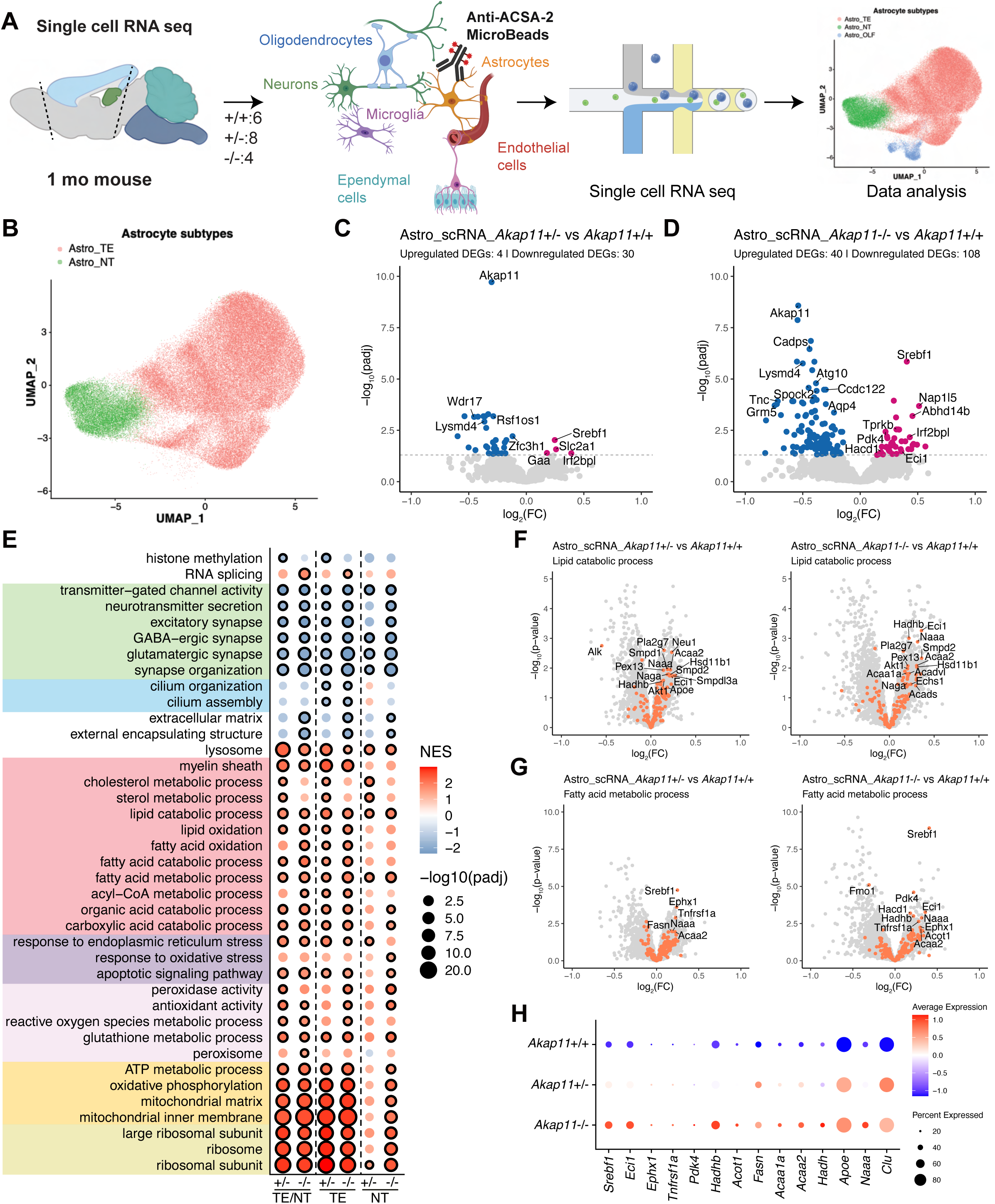
ScRNA-seq of astrocytes from *Akap11*^+/+^, *Akap11*^+/-^, and *Akap11*^-/-^ mice reveals transcriptomic changes linked to lipid dysregulation. **(A)** Overview of the scRNA-seq data generation and analysis of astrocytes from 1-month-old *Akap11*^+/+^, *Akap11*^+/-^, and *Akap11*^-/-^ mice. Cerebral hemispheres were dissected and dissociated, followed by astrocyte isolation using magnetic-activated cell sorting (MACS) targeting the astrocyte-specific surface antigen ACSA2, and single cell barcoding for scRNA-seq. The number of mice used for each genotype is indicated. **(B)** Uniform manifold approximation and projection (UMAP) plot showing two distinct astrocyte subtypes identified by scRNA-seq: telencephalon astrocytes (Astro_TE) and non-telencephalon astrocytes (Astro_NT). **(C and D)** Volcano plots displaying significantly altered genes (FDR < 0.05) in *Akap11*^+/-^ (**C**) and *Akap11*^-/-^ (**D**) astrocytes compared to *Akap11*^+/+^ controls. For differential expression analysis, the astro-TE and astro-NT subtypes were combined. The horizontal dotted line indicates the significance threshold (FDR = 0.05). Genes upregulated in mutants are shown in red; downregulated genes are shown in blue. **(E)** Bubble plot of GSEA results highlighting changes in select molecular pathways. Significant changes (FDR < 0.05) have black outer circles. *Akap11*^+/-^ and *Akap11*^-/-^ astrocytes were compared to *Akap11*^+/+^. **(F and G)** Volcano plots highlighting transcriptomic changes in *Akap11*^+/-^ and *Akap11*^-/-^ mouse astrocytes compared to *Akap11*^+/+^ controls. (**F**) highlights genes associated with the lipid catabolic process, and (**G**) highlights those involved in the fatty acid metabolic process. Orange dots indicate genes within the respective gene sets. **(H)** Dot plot showing relative expression levels for representative genes associated with lipid catabolic and fatty acid metabolic processes.

Differential gene expression analysis using a pseudobulk approach revealed that *Srebf1* is the most significantly upregulated transcript in both *Akap11*^+/-^ and *Akap11*^-/-^ astrocytes compared to *Akap11*^+/+^ controls (Figures, 2C and 2D). *Srebf1* encodes the key transcription factor that drives the expression of genes involved in cholesterol and lipid biosynthesis^39^. Additionally, *Eci1*, essential for fatty acid β-oxidation; *Hacd1*, required for long-chain fatty acid elongation; and *Pdk4*, a critical regulator of glucose and lipid metabolism, were significantly upregulated in *Akap11*^-/-^ astrocytes (Figure 2D). *Pdk4* upregulation indicates a metabolic shift towards enhanced fatty acid utilization as an energy source, serving as a sensitive marker of increased fatty acid oxidation^40^.

To explore the biological processes affected in *Akap11*^+/-^ and *Akap11*^-/-^ mouse astrocytes, we performed GSEA on scRNA-seq from individual astrocyte subtypes (Astro-TE and Astro-NT), as well as pooled TE and NT astrocytes (Astro-TE/NT). Astro-TE and Astro-NT subtypes displayed similar GSEA enrichment patterns, and their pathway changes largely mirrorred those observed in cultured astrocytes, except for synapse related terms (Figures 2E and 1B). This suggests that RNA-seq data from cultured astrocytes generally resembles our *in vivo* findings, particularly in the metabolism/mitochondria/ribosome-related pathways. For instance, mitochondrial/OXPHOS and ribosome-related Gene Ontology (GO) terms were significantly upregulated in both *Akap11*^+/-^ and *Akap11*^-/-^ astrocytes. Lipid-related pathways, including those involved in cholesterol/sterol metabolism, fatty acid/lipid catabolism, and fatty acid/lipid oxidation, were also significantly upregulated, particularly in the Astro-TE subtype. Additionally, pathways associated with peroxisomes, ER stress, oxidative stress, ROS, peroxidative activity, and apoptotic signaling, which are closely linked to lipid metabolism or mitochondrial respiration, were upregulated. In line with observations in cultured astrocytes, cilium-related terms were downregulated in GSEA, specifically in the Astro-TE subtype (Figure 2E). Interestingly, however, GO terms related to synapses and neurotransmitters were significantly downregulated in astrocytes acutely isolated from the brain, which is opposite to the upregulation observed in cultured astrocytes (Figures 2E and 1B). This discrepancy might result from *Akap11* loss of function affecting astrocyte-neuron interactions *in vivo*, which is not replicated in primary astrocyte cultures that lack neurons.

Which specific genes are upregulated in lipid catabolic processes and fatty acid metabolism in astrocytes acutely isolated from brain? In addition to above highlighted genes like *Srebf1, Eci1, Hacd1,* and *Pdk4*, we observed upregulation of *Apoe* and *Clu* (Clusterin), also known as *apolipoprotein J* (*ApoJ*), in both *Akap11*^+/-^ and *Akap11*^-/-^ astrocytes (Figures 2F and 2H). *Apoe* and *Clu* are recognized as genetic risk factors for Late-Onset Alzheimer’s Disease^41^ and are among the principal apolipoproteins responsible for cholesterol transport in the brain^42^. We also detected increased expression of *Smpd1* and *Smpd2*, which encode acid sphingomyelinases (ASM) that catalyze the conversion of sphingomyelin to ceramide (Figure 2F), as well as *ceramide synthase 2* (*Cers2*), which is responsible for the synthesis of very long-chain ceramides (Table S3). Additional upregulated genes involved in fatty acid oxidation included *Echs1*, *Eci1*, *Hadh*, *Hadhb*, *Acaa2*, and *Acaa1a*, along with *Fasn*, which directly catalyzes the de novo biosynthesis of long-chain saturated fatty acids (Figures, 2F-2H and S3F). Importantly, many of these genes, such as *Apoe*, *Acaa1a*, *Acaa2*, *Acads*, *Acadvl*, *Agt*, *Cyb5a*, *Ech1*, *Etfb*, *Eci1*, *Ephx1*, *Fasn*, *Hadh*, and *Hadhb*, also showed an upregulation in cultured astrocytes, underscoring the consistency between our *in vitro* and *in vivo* findings (Figure S3F and Table S3). A transcriptome-wide comparison further revealed that log_2_FC values of individual genes were positively correlated between cultured astrocytes and *in vivo* mouse astrocytes (Figure S3G and S3H).

### *Akap11* deficiency causes accumulation of cholesteryl esters, triacylglycerols, ceramides, and glycerophospholipids in cultured astrocytes

Our RNA-seq and proteomic analyses revealed significant upregulation of genes and proteins associated with lipid metabolism in *Akap11* deficient astrocytes. Specifically, we observed increased expression of genes related to cholesterol transport (*Apoe, Clu, Abca1*) and uptake (*Lrp1, Lrp5, Vldlr*), as well as genes involved in cholesteryl ester synthesis (*Acat2, Acat3*) and ceramide synthesis (*Smpd1, Smpd2, Cers2*). Additionally, genes associated with fatty acid synthesis (*Fasn*), transport (*Mfsd2a, Cpt1, Fabp3*), and oxidation (*Ech1, Eci1, Hadh, Hadhb, Acaa2, Acaa1a*) were increased. Notably, there was enhanced RNA expression of transcription factor *Srebf1*, a key regulator of cholesterol biosynthesis and lipid homeostasis (Figure 2C and 2D). To directly measure changes in levels of lipids in unbiased fashion, we conducted liquid chromatography-mass spectrometry (LC-MS)-based lipidomic and metabolomic profiling on cultured astrocytes from *Akap11*^+/+^, *Akap11*^+/-^, and *Akap11*^-/-^ mice (Figure 3A). This analysis identified a total of 780 annotated metabolites (Table S4), with 143 upregulated and 8 downregulated metabolites in cultured *Akap11*^-/-^ astrocytes, and 34 upregulated and 3 downregulated metabolites in cultured *Akap11*^+/-^ astrocytes (nominal p < 0.05) (Figure S4A). Among these, 71 lipid species were significantly increased in *Akap11*^-/-^ astrocytes compared to *Akap11*^+/+^ controls, while 7 lipid species were decreased (Figure 3B; Table S4). Individual metabolite log₂FC values were well correlated across genotypes (r = 0.67; Figure 3C).

**Figure 3.**
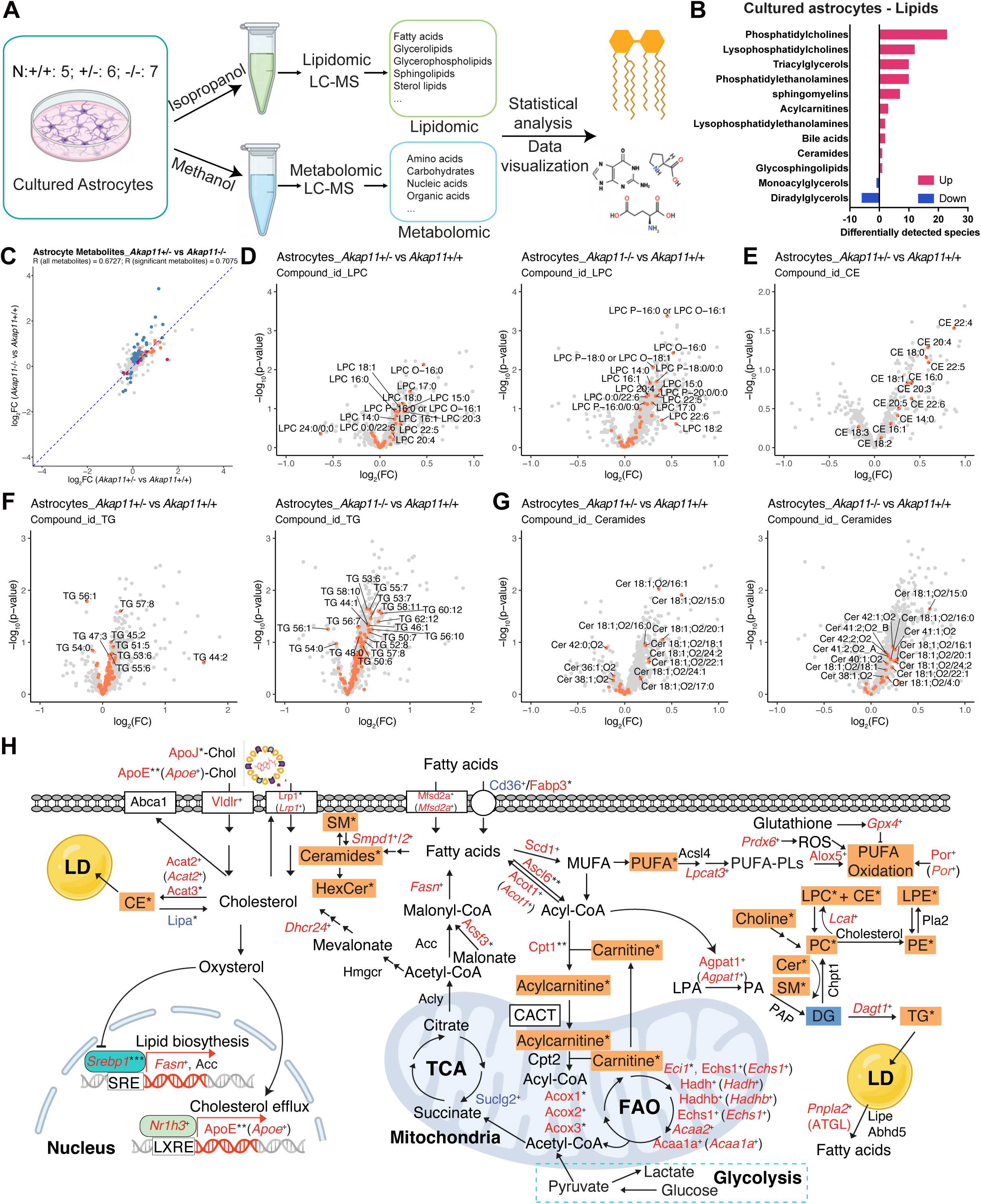
Loss of *Akap11* leads to excessive lipid accumulation in cultured astrocytes. **(A)** Schematic overview of lipidomic and metabolomic analyses performed on cultured astrocytes isolated from *Akap11*^+/+^, *Akap11*^+/-^, and *Akap11*^-/-^ pups prepared as described in Figure 1A. The number of mice used for astrocytes isolation is indicated for each genotype. **(B)** Bar chart showing the number of differentially detected lipid species (p < 0.05) for each lipid class in *Akap11*^-/-^ cultured astrocytes compared to *Akap11*^+/+^. **(C)** Correlation of metabolites between *Akap11*^+/-^ vs *Akap11*^+/+^ and *Akap11*^-/-^ vs *Akap11*^+/+^ cultured astrocytes. Metabolites significantly altered (p < 0.05) in the *Akap11*^+/-^ comparison are labeled in red, in the *Akap11*^-/-^ comparison in blue, and in both comparisons in orange. **(D-G)** Volcano plots showing changes in specific lipid species in *Akap11*^+/-^ and *Akap11*^-/-^ astrocytes compared to *Akap11*^+/+^ astrocytes, focusing on lysophosphatidylcholines (LPC, **D**), cholesteryl esters (CE, **E**), triacylglycerols (TG, **F**), and ceramides (**G**). CE data is derived from an independent experiment where astrocytes were grown in cholesterol-free medium. Orange dots indicate lipids belonging to the respective lipid class. **(H)** Integrated analysis of transcriptomic, proteomic, and metabolomic data from *Akap11* mutant astrocytes. Differentially expressed genes (italicized) and proteins are color-coded (red: increased; blue: decreased) with statistical significance denoted as follows: p-value < 0.05, “+”; padj ≤ 0.1, “.”; padj < 0.05, “*”; padj < 0.01, “**”; padj < 0.001, “***”. Changes in metabolite abundance are indicated by colored rectangles (orange: increased; blue: decreased; p-value < 0.05, “*”).

The most significantly increased lipid species included phosphatidylcholines (PC), lysophosphatidylcholines (LPC), triacylglycerols (TG), phosphatidylethanolamines (PE), sphingomyelins (SM), acylcarnitines (CAR), lysophosphatidylethanolamines (LPE), and ceramides. The increase in LPC, PC, LPE, and PE levels occurred in a gene-dose dependent fashion in *Akap11*^+/-^ and *Akap11*^-/-^ astrocytes (Figures 3D and S4B-S4D). We also observed broad remodeling of ether-linked phospholipids. Specifically, levels of alkyl ether-linked (O-series) and plasmalogen-type (P-series) lipids were elevated across multiple classes, including LPC (LPC O-/P-), PC (PC O-/P-), and PE (PE O-/P-). These ether lipids play critical roles in membrane structure and antioxidant defense and are often upregulated in response to oxidative and metabolic stress.

Consistent with the upregulation of genes involved in cholesterol transport, uptake, and esterification, *Akap11*-deficient astrocytes exhibited elevated levels of CE (Figures 3E and S4E). We also observed a rise in lipid droplet TG species, which are typically formed via *de novo* lipogenesis through fatty acid synthesis and subsequent esterification into TGs^43^. The increased expression of *Fasn* and *Dgat1*, key enzymes in this process, supports this mechanism (Figure 3F). In parallel, genes associated with ceramide synthesis (*Smpd1*, *Smpd2*, *Cers2*) were upregulated, matching the observed increase in ceramide levels (Figure 3G). Additionally, sphingomyelin (SM) species were elevated, further indicating dysregulation of sphingolipid metabolism in *Akap11*-deficient astrocytes (Figure S4F).

Beyond lipid synthesis and storage, we also detected changes related to lipid peroxidation. Analysis of lipid peroxidation byproducts revealed elevated levels of 13-Hydroxyoctadecadienoic acid (13-HODE), a peroxidation product derived from linoleic acid, in *Akap11* mutant astrocytes (Figure S4G). This increase aligns with our transcriptomic data, which suggests heightened oxidative stress in these cells (Figures 1B and 2E). Supporting this notion, there was increased expression of genes involved in iron-mediated lipid peroxidation (*Cdo1*, *Ftl1*, *Fth1*, *Plod1*, *Iscu*, *Gpx4*, *Slc3a2*, *Lpcat3*, *Coq10b*, *Por* and *Prdx1-Prdx6*) (Figure S4H). Intriguingly, we also observed a concurrent upregulation of fatty acid oxidation genes, which correlated with higher acylcarnitine levels, a key intermediate in β-oxidation responsible for transporting long-chain fatty acids into mitochondria^44^ (Figure S4I). Figure 3H shows an integrated view of how *Akap11* deficiency reshapes cellular metabolism in astrocytes, based on transcriptomic, proteomic, lipidomic, and metabolomic data (Figure 3H). Collectively, our findings demonstrate that *Akap11* deficiency causes widespread lipid metabolic dysregulation in cultured astrocytes, leading to the accumulation of CE, TG, ceramides, and glycerophospholipids alongside transcriptional upregulation of cholesterol transport, fatty acid synthesis, and oxidation genes.

### *Akap11* deficiency causes cholesteryl esters, triacylglycerols, ceramides, and glycerophospholipids accumulation in the brain

Our observation of profound lipid dysregulation in cultured astrocytes prompted us to ask whether similar alterations occur in *Akap11*-deficient brains. To explore this, we performed lipidomic and metabolomic analyses on cerebral hemispheres from 1-month-old *Akap11*^+/+^, *Akap11*^+/-^, and *Akap11*^-/-^ mice (Figure 4A). This analysis identified a total of 832 annotated metabolites (Table S5), with 163 upregulated and 49 downregulated metabolites in *Akap11*^-/-^ mice, and 61 upregulated and 3 downregulated metabolites in *Akap11*^+/-^ mice (nominal p < 0.05) (Figure 4C). Among the 832 metabolites, 115 lipid species were significantly increased in *Akap11*^-/-^ mice compared to *Akap11*^+/+^ controls, while 22 species were decreased (Figure 4B; Table S5). Consistent with our astrocyte findings, PC, LPC, LPE, ceramides, SM, CE, and TG were all increased in *Akap11* mutant brains (Figures, 4B and 4C). Notably, log₂FC in individual metabolite levels were strongly correlated across genotypes (r = 0.79; Figure S5B), supporting a gene-dose–dependent effect of *Akap11* loss on brain lipid homeostasis.

**Figure 4.**
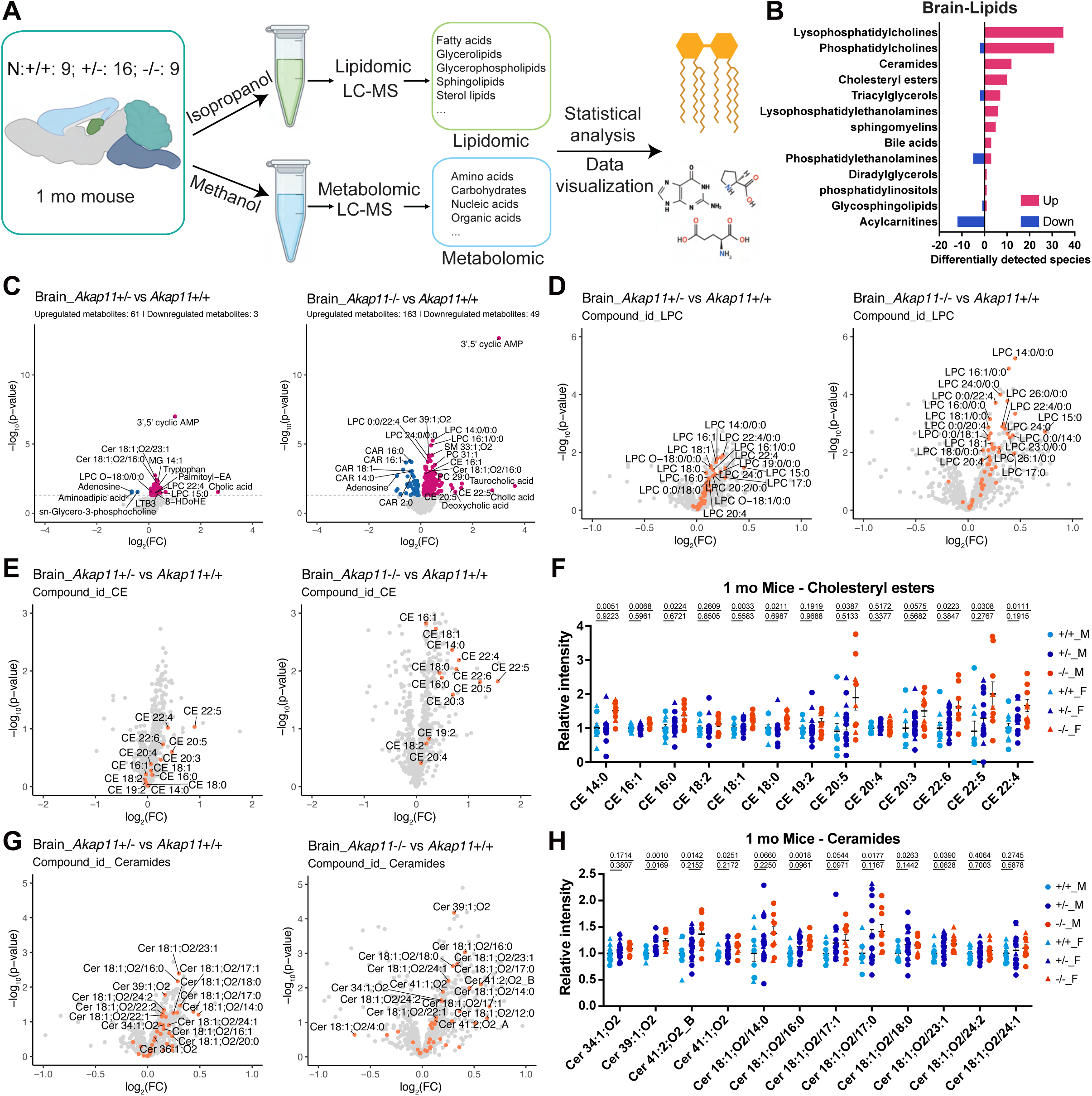
Lipid accumulation in *Akap11* mutant brains. **(A)** Schematic overview of lipidomic and metabolomic analyses of cerebral hemispheres from 1-month-old *Akap11*^+/+^, *Akap11*^+/-^, and *Akap11*^-/-^ mice. Sample sizes (number of mice) per genotype are indicated. **(B)** Bar chart illustrating the number of significantly altered lipid species (p < 0.05) across lipid classes in *Akap11*^-/-^ mice compared to *Akap11*^+/+^ controls. **(C)** Volcano plots showing significantly (p < 0.05) increased (red) or decreased (blue) lipid species and metabolites in *Akap11*^+/-^ mice (left) and *Akap11*^-/-^ mice (right) compared to *Akap11*^+/+^ controls. The horizontal dotted line indicates the significance threshold (p = 0.05). **(D, E, G)** Volcano plots highlighting altered lipid species, specifically lysophosphatidylcholines (LPC, **D**), cholesterol esters (CE, **E**), and ceramides (**G**), in *Akap11*^+/-^ or *Akap11*^-/-^ mice compared to *Akap11*^+/+^ mice. Orange dots indicate lipids belonging to the respective lipid class. **(F)** Relative abundance of all detected CE species in brain extracts from 1-month-old *Akap11*^+/+^, *Akap11*^+/-^, and *Akap11*^-/-^ mice. **(H)** Relative abundance of representative ceramide species in brain extracts from 1-month-old *Akap11*^+/+^, *Akap11*^+/-^, and *Akap11*^-/-^ mice.

An unbiased network analysis integrating metabolomic and lipidomic data with RNA-seq (scRNA-seq) data using the GAM (’genes and metabolites’) tool^45^ revealed extensive alterations in lipidomic and metabolic pathways in *Akap11*^-/-^ mice compared to *Akap11*^+/+^ mice (Figure S5A). Specifically, *Akap11*^-/-^ mice exhibited: (1) an accumulation of lipid species TG, CE, PC, and LPC; (2) a marked reduction in key intermediates of the pentose phosphate pathway, such as 6-phosphogluconate and ribose 5-phosphate; (3) decreased levels of glycolysis-related metabolites, including glucose, fructose 1,6-bisphosphate, and pyruvate, as well as a reduction in anaerobic glycolysis products like lactate; (4) A decrease in antioxidants glutathione (GSH) and glutathione disulfide (GSSG); (5) elevated levels of catabolic products of amino acids and their derivatives, cystine, methionine, and S-adenosylhomocysteine; and (6) decreased levels of metabolites associated with fatty acid oxidation, such as palmitoylcarnitine and acetylcarnitine (Figures 4C and S5A).

We found that PC, LPC, and LPE levels increased in a gene dose-dependent manner in *Akap11*^+/-^ and *Akap11*^-/-^ brain, similar to cultured astrocytes (see above), with LPC and LPE showing the most pronounced elevations compared to *Akap11*^+/+^ controls (Figures 4D and S5C-S5F). Similarly, CEs were significantly higher in *Akap11*^-/-^ mice and were intermediate in *Akap11*^+/-^ mice compared to *Akap11*^+/+^ controls (Figures 4E and 4F). Notably, specific CE species (CE22:4, CE18:0, and CE22:5) that were elevated in *Akap11* mutant astrocytes (Figure 3E) showed parallel increases in the mutant brain. While CE levels rose, overall brain cholesterol levels remained unchanged, and cholesterol precursors lanosterol and 14-Demethyllanosterol were reduced (Figure S5G). TGs also showed a gene dose-dependent increase in *Akap11*^+/-^ and *Akap11*^-/-^ mice, though the increase in *Akap11*^+/-^ mice was not statistically significant (Figures S5H and S5I). In addition, ceramides, lipids linked to apoptosis, oxidative stress and inflammation, and their glucosylated form, glucosylceramide (GlcCer), were notably higher in both *Akap11*^+/-^ and *Akap11*^-/-^ mice (Figures 4G, 4H and S6A). Sphingomyelin (SM), a critical component of the myelin sheath, also showed a slight increase in these mutant mice (Figure S6B). Interestingly, elevated ceramide levels have been observed in prefrontal cortex tissue from individuals with either SCZ or BD^46^, as well as in the serum or plasma of patients with major depressive disorder, linking these lipid alterations to a broader range of psychiatric conditions^47,48^.

Unlike cultured astrocytes, where acylcarnitine levels were increased, *Akap11*^-/-^ brains exhibited reduced acylcarnitine levels, while *Akap11*^+/-^ brains showed no significant change (Figure S6C). Acylcarnitine helps transport long-chain fatty acids into mitochondria for β-oxidation, so its decrease likely leads to fatty acid buildup in the cytoplasm and less fuel for β-oxidation. Consistent with this, we detected significant increases in several long-chain fatty acids in *Akap11* mutant mice, including margaric acid, alpha-linolenic acid, palmitoleic acid, and myristic acid (Figure S6D). Polyunsaturated fatty acids (PUFAs), such as docosahexaenoic acid (DHA), arachidonic acid (ARA), and linolenic acid, are particularly susceptible to peroxidation, which, when extensive, damages membranes and triggers ferroptosis. In *Akap11*^+/-^ but not *Akap11*^-/-^ mice, we observed elevated levels of oxidative PUFA byproducts, including hydroxyoctadecadienoic acids (HODEs, from linoleic acid), hydroxyeicosatetraenoic acids (HETEs, from arachidonic acid), and hydroxydocosahexaenoic acids (HDoHE, from DHA) (Figure S6E). These changes align with increased levels of lipoxygenase (ALOX) proteins in the brain (Figure S6F), which catalyze PUFA peroxidation to form lipid peroxyl radicals, and with decreased glutathione (GSH) (Figures S6F and S5A), an antioxidant whose reduction promotes lipid peroxidation^49^. Interestingly, oxidative stress and lipid peroxidation may contribute to the pathology of SCZ, BD, and other psychiatric disorders^50–52^. We summarized the main lipid metabolic pathways and the related RNA expression changes (based on bulk RNA-seq data from the hippocampus (Hipp) and prefrontal cortex (PFC)^27^) in *Akap11* mutant mice (Figure S6G).

### *Akap11* deficiency elevates PKA subunits, 3’,5’ cyclic AMP, and PKA activity

What are the mechanisms by which AKAP11 regulates lipid metabolism in astrocytes? The cAMP-PKA signaling pathway is a well-established regulator of lipid metabolism, particularly in adipocytes, where PKA phosphorylates lipases on the surfaces of lipid droplets, promoting lipolysis^53–57^. However, in non-adipose tissues, the effects of cAMP-PKA signaling on lipid droplets can vary. For instance, cAMP or PKA activation has been shown to increase lipid droplet accumulation in astrocytes^58^, hepatocytes^59^, breast cancer cells^60^, and Rat sertoli cells^61^. Additionally, PKA directly phosphorylates SREBP1, a transcription factor critical for fatty acid and cholesterol biosynthesis, modulating its transcriptional activity^62^.

Our proteomics analysis of *Akap11* deficient astrocytes revealed a significant elevation of PKA subunits, including PKA Cα, PKA Cβ (encoded by *Prkaca* and *Prkacb*), PKA R1α, PKA R1β (encoded by *Prkar1a* and *Prkar1b*), and PRKX (encoded by *Prkx*) in both *Akap11*^+/-^ and *Akap11*^-/-^ astrocytes (Fig. 5A). Marked accumulation of PKA proteins also occurs in the brain of *Akap11* mutant mice^27,63^. There were no significant changes in the mRNA levels of these PKA subunits in mutant astrocytes (Figure S7A), suggesting that AKAP11 predominantly regulates PKA protein abundance at the post-transcriptional level, which aligns with previous findings that AKAP11 mediates PKA subunits turnover through autophagy in human cell lines and human pluripotent stem cell (PSC)-induced neurons^21,22,64^. Western blot analysis confirmed these proteomics results: in *Akap11*^-/-^ cultured astrocytes, PKA R1-α levels increased ∼10-fold and PKA C-α levels increased approximately 4-fold. In *Akap11*^+/-^ cultured astrocytes, PKA R1-α levels rose by 1.5- to 2-fold and PKA C-α levels increased by 1.5-fold (Figures 5B and 5C). Similar patterns were observed in *Akap11*^-/-^ embryonic cortical neurons, where PKA R1-α increased by 7-fold and PKA C-α by 1.7-fold. Although the PKA increases in *Akap11*^+/-^ samples did not reach statistical significance, a similar trend was observed (Figures S7B and S7C). The elevation of PKA subunits was also seen in cultured rat hippocampal neurons in which AKAP11 expression was knocked down by siRNA and in cultured HEK293T cells in which *AKAP11* was disrupted by CRISPR-Cas9 editing (Figures S7D and S7E). Thus, the marked elevation of PKA subunits is a consistent response to *Akap11* deficiency across different cell types and different species.

**Figure 5.**
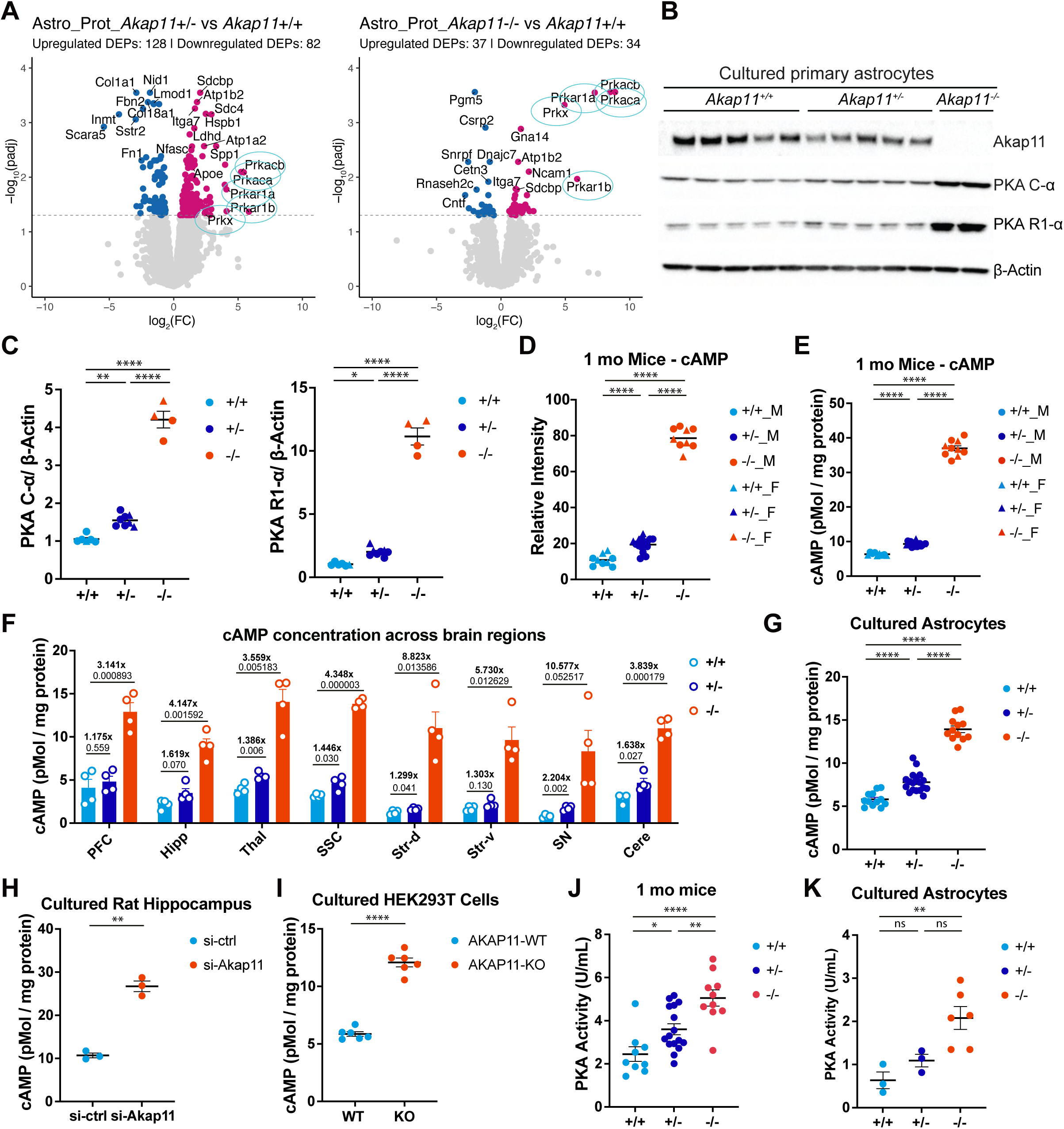
*Akap11* deficiency increases PKA subunits, cAMP, and PKA activity. **(A)** Volcano plots showing significantly altered proteins (FDR < 0.05) in *Akap11*^+/-^ and *Akap11*^-/-^ cultured astrocytes compared to *Akap11*^+/+^. PKA subunits are highlighted with cyan-blue circles. Log_2_FC values resulting from the linear model applied to the dataset after Median-MAD normalization (see Methods for details). The horizontal dotted line indicates the significance threshold (FDR = 0.05). Upregulated and downregulated proteins are shown in red and blue, respectively. **(B)** Immunoblot analysis of protein lysates from *Akap11*^+/+^, *Akap11*^+/-^, and *Akap11*^-/-^ astrocytes using the indicated antibodies. **(C)** Quantification of immunoblot results from (**B**) with protein levels normalized to β-Actin. Data are presented as mean ± SEM (n ≥ 4 mice/genotype). Statistical significance was determined by one-way ANOVA followed by Tukey’s multiple comparisons test. *p<0.05; **p<0.01; ***p<0.001; ****p<0.0001; ns, not significant. **(D)** Relative cAMP level in brain extracts from 1-month-old *Akap11*^+/+^, *Akap11*^+/-^, and *Akap11*^-/-^ mice, as determined by LC-MS. Statistical significance was performed using one-way ANOVA followed by Tukey’s multiple comparisons test. ****p<0.0001. **(E and F)** Quantification of cAMP level in cerebral hemisphere brain extracts (**E**) and specific brain regions (**F**) from 1-month-old *Akap11*^+/+^, *Akap11*^+/-^, and *Akap11*^-/-^ mice by LANCE Ultra cAMP assays. Each dot represents one mouse. Statistical significance was determined by one-way ANOVA followed by Tukey’s multiple comparisons test for (**E**) and multiple unpaired t test for (**F**). ****p<0.0001. **(G-I)** Quantification of cAMP level in primary cultured astrocytes from *Akap11*^+/+^, *Akap11*^+/-^, and *Akap11*^-/-^ mice (**G**; each dot represents an individual cell culture, N mice: *Akap11*^+/+^ = 2, *Akap11*^+/-^ = 6, *Akap11*^-/-^ = 4), cultured rat hippocampal neurons (**H**), and cultured HEK293T cells (**I**) using LANCE Ultra cAMP assays. Statistical analysis was performed using one-way ANOVA followed by Tukey’s multiple comparisons test for (**G**), and Welch’s t-tests for (**H**) and (**I**). *p<0.05; **p<0.01; ***p<0.001; ****p<0.0001; ns, not significant. **(J and K)** Quantification of PKA activity in 1-month-old mice (**J**; each dot represents an individual mouse, N mice: *Akap11*^+/+^ = 9, *Akap11*^+/-^ = 16, *Akap11*^-/-^ = 10) and cultured astrocytes (**K**; each dot represents cells from one mouse) from *Akap11*^+/+^, *Akap11*^+/-^, and *Akap11*^-/-^ mice using a PKA Colorimetric Activity Kit. Statistical significance was determined by one-way ANOVA followed by Tukey’s multiple comparisons test. *p<0.05; **p<0.01; ***p<0.001; ****p<0.0001; ns, not significant.

To further explore the mechanism of AKAP11-mediated PKA autophagy, we induced autophagy in cultured astrocytes through nutrient starvation and observed a reduction of PKA R1-α and PKA C-α levels in *Akap11*^+/+^ astrocytes, a partial reduction in *Akap11*^+/-^ astrocytes, and no reduction in *Akap11*^-/-^ astrocytes (Figures S7F and S7G). Under nutrient starvation and Baf A1 (a specific vacuolar H^+^ ATPase inhibitor) treatment, PKA R1-α formed puncta that colocalized with the endogenous autophagosome marker P62 (Figure S7H). Notably, the formation of these puncta was decreased in *Akap11*^+/-^ and abolished in *Akap11*^-/-^ astrocytes under the same conditions, suggesting an essential role of AKAP11 in recruiting PKA R1-α to autophagosomes (Figure S7H). Together, these findings indicate that AKAP11 mediates the autophagic degradation of PKA subunits in astrocytes and neurons and is required for suppression of PKA levels during nutrient deprivation.

PKA is the major effector for the second-messenger molecule cAMP. Remarkably, we found that cAMP was highly elevated in *Akap11* mutant mice by unbiased metabolomics profiling (Figures 4C and 5D). Specifically, cAMP levels increased approximately 1.5- to 2-fold in total brain of *Akap11*^+/-^ mice, and about 7- to 8-fold in *Akap11*^-/-^ mice, with no sex differences (Figure 5D). To validate this finding, we measured cAMP using a time-resolved fluorescence resonance energy transfer (TR-FRET)-based immunoassay. In wild-type (WT) brain tissue, baseline cAMP levels were 6.5 pmol/mg protein. Consistent with the metabolomic data, cAMP levels increased 1.5-fold in *Akap11*^+/-^ mice and 6-fold in *Akap11*^-/-^ mice compared to Akap11^+/+^ mice (Figure 5E). This remarkable elevation of cAMP in *Akap11* mutants was observed across all examined brain regions: the prefrontal cortex (PFC), hippocampus (Hipp), thalamus (Thal), somatosensory cortex (SSC), dorsal striatum (Str-d), ventral striatum (Str-v), substantia nigra (SN), and cerebellum (Cere). The largest fold increases in *Akap11* mutants versus WT were found in the dorsal striatum and substantia nigra (Figure 5F). We also observed elevated cAMP in cultured cerebral astrocytes derived from *Akap11*^+/+^, *Akap11*^+/-^, and *Akap11*^-/-^ postnatal mouse pups (P1-P4) (Figure 5G), rat hippocampus cultures treated with *Akap11* siRNA (Figure 5H), and human HEK293T cells in which *AKAP11* has been knocked out by CRISPR-Cas9 (Figure 5I), suggesting that cAMP elevation is a universal consequence of *Akap11* deficiency. Next, we examined PKA kinase activity in *Akap11* mutant mice and cultured astrocytes using a commercial PKA colorimetric activity kit. PKA activity was found to increase in a gene dose-dependent manner in *Akap11* mutant mice and cultured astrocytes (Figures 5J and 5K).

We hypothesized that AKAP11 regulates lipid metabolism in astrocytes by modulating cAMP levels, PKA activity, or downstream phosphorylation targets (Figure S8D). To identify specific phosphorylation targets influenced by *Akap11* deletion in astrocytes, we conducted quantitative phosphoproteomic profiling using tandem mass tag (TMT) labeling in *Akap11*^+/+^, *Akap11*^+/-^, and *Akap11*^-/-^ astrocytes (Figures S8A and S8B; Table S2). This analysis identified significant phosphorylation changes (FDR < 0.1) at multiple sites on key proteins involved in lipid metabolism, including Pcyt1a, Osbpl3, Lpar1, Smpd3, Plcl2, Cp (ceruloplasmin), Acot11, Spp1, and Pitpnm3 (Figures S8C; Table S2). These results support the idea that dysregulated cAMP-PKA signaling contributes to altered lipid metabolism in *Akap11*-deficient astrocytes.

### AKAP11 interacts with VAPs via FFAT motif

Neutral lipids like TGs and cholesteryl esters CEs are mainly stored in lipid droplets, which are surrounded by a single phospholipid monolayer. To directly visualize lipid accumulation in *Akap11* mutant astrocytes, we used BODIPY™ 493/503, a neutral lipid-specific dye. The staining showed a significant increase of lipid droplet numbers and intensity, particularly in *Akap11*^+/-^ astrocytes (Figures S9A and S9B).

Originating from the ER, lipid droplets are highly dynamic organelles that can interact with many other cellular components through membrane contact sites. Their formation, degradation, and interactions with other organelles are closely tied to cellular metabolism and help buffer cells against excess or toxic lipid levels^65,66^. We recently showed that AKAP11 interacts with vesicle-associated membrane protein-associated proteins VAP-A and VAP-B in the brain^27^. VAPs primarily function to tether the ER to various other organelles and regulate lipid transport and metabolism^67–69^. They are also known to interact with multiple autophagy proteins and modulate autophagosome biogenesis^70,71^. Based on this, we hypothesize that the AKAP11-VAP complex plays a key role in AKAP11-mediated lipid metabolism (Figure S8D).

We verified the interaction between AKAP11 and VAPs using affinity purification coupled with mass spectrometry (AP-MS) of EGFP-tagged AKAP11 in HEK293T cells. This analysis identified 71 significantly enriched proteins (P < 0.05) from 2,294 detected co-immunoprecipitated proteins across two independent experiments (Figure S9C; Table S6). Among the 71 enriched proteins, AKAP11 was the most enriched in the affinity purification, as expected, alongside seven previously known AKAP11 interactors: PKA subunits (PKA R1-α (PRKAR1A), PKA C-α (PRKACA), and PKA C-β (PRKACB)), IQGAP1, IQGAP3, VAP-A, and VAP-B^19,20,23,72^. Additionally, PKA R2-α (PRKAR2A) was significantly enriched in one of the co-IP experiments (Table S6). Beyond these known interactors, we identified several novel AKAP11-binding partners. These include UFD1-VCP-NPLOC4 ternary complex, 26S proteasome subunits, and E3 ligase HUWE1 and its scaffold CUL2, which are proteins involved in endoplasmic reticulum-associated degradation (ERAD), the centrosomal protein CEP170, which supports cilia function through interaction with dynein-2^73^, and ATAD3A, ATAD3B, FAR1, ABCD3, and ACOT8 which are proteins involved in lipid metabolism (Figure S9C; Table S6).

We validated several of these interactors through co-IP-immunoblot experiments from mouse brain^27^ and HEK293T cells, including PKA R1α, PKA Cα, VAP-A, and VAP-B (Figure S9D). Additionally, AKAP11 was found to associate with mitochondrial proteins (Figure S9D), especially ATAD3A, which spans the inner and outer mitochondrial membranes and is required for cholesterol transport^74^. Furthermore, AKAP11 binds to ABCD3/PMP70 (Figure S9D), a peroxisomal ATP-binding cassette (ABC) transporter essential for the transport of fatty acids and fatty acyl-CoAs from the cytosol into peroxisomes^75^. Notably, *Abcd3* knockout mice display perturbations in hepatic cholesterol synthesis^76^.

VAPs function to anchor soluble cytosolic proteins to the cytoplasmic surface of the ER. This anchoring is often mediated by the FFAT motif (two phenylalanines in an acidic tract) present in VAP-associated proteins, which facilitates direct interaction with the major sperm protein (MSP) domain of VAPs^72,77,78^. Intriguingly, AKAP11 also contains a putative FFAT motif^79^ spanning amino acids 354-360, which is conserved among other VAP-associated proteins (Figure S9E). To explore this interaction further, we employed AlphaFold 3 to predict the structure of AKAP11-VAP complexes^80^. In this model, the AKAP11 FFAT motif “EFFDSFD” lies at the interface between AKAP11 and both VAP-A and VAP-B (Figure S9F). Structure prediction suggested that E354, D357, and D360 are the key residues within FFAT at the AKAP11-VAP interface (Figure S9F). To experimentally validate the role of these residues, we conducted a series of co-IP experiments with point mutants of AKAP11. EGFP-tagged AKAP11 with individual mutations of these residues decreased VAP interactions, while the triple mutant E354AD357AD360A almost abolished VAP binding (Figure S9G). Importantly, these mutations did not affect AKAP11’s interaction with PKA subunits or the peroxisomal membrane protein PMP70 (Figure S9G). Collectively, these results indicate that the FFAT motif in AKAP11 mediates its interaction with VAPs. Future studies will test whether this AKAP11-VAP interaction is necessary for AKAP11’s role in regulating lipid metabolism (Figure S8D).

### *Akap11* deficiency in astrocytes increases neuronal activity of co-cultured neurons

Astrocytes play a critical role in neuronal function by regulating neurotransmitter levels, providing metabolic and energy support, modulating synaptic activity and plasticity, maintaining the blood-brain barrier, and neurorepair^81–85^. They are also considered the primary source of cholesterol synthesis in the brain, supplying essential sterols to neurons, which are crucial for synapse formation^42,83,86^. The coupling of lipid metabolism between astrocytes and neurons has been reported as important for maintaining proper neuronal activity^87–89^. To study how *Akap11* loss of function in astrocytes affects neurons, we co-cultured mouse astrocytes (*Akap11*^+/+^, *Akap11*^+/-^, and *Akap11*^-/-^) with human induced pluripotent stem cell (hiPSC)-derived neurons. Using hiPSC-neurons avoids glial contamination, and we can distinguish astrocyte (mouse) from neuron (human) RNAs by RNA sequencing, which we used to analyze the molecular effects of *Akap11* mutant astrocytes on human gene expression in iPSC neurons.

We performed RNA-seq on cocultures collected on DIV28 and DIV42 (Figure 6A), time points at which neurons co-cultured with glial cells exhibit well-defined morphology and robust electrophysiological activity^90,91^. We confirmed that RNA from human neurons could be reliably distinguished from those originating from mouse glial cells by alignment of RNA-seq reads to human or mouse references (Figure S10A). Principal component analysis (PCA) of the RNA expression profiles revealed that co-culturing with astrocytes induced major transcriptional changes in neurons at both days DIV28 and DIV42 (Figure S10B).

**Figure 6.**
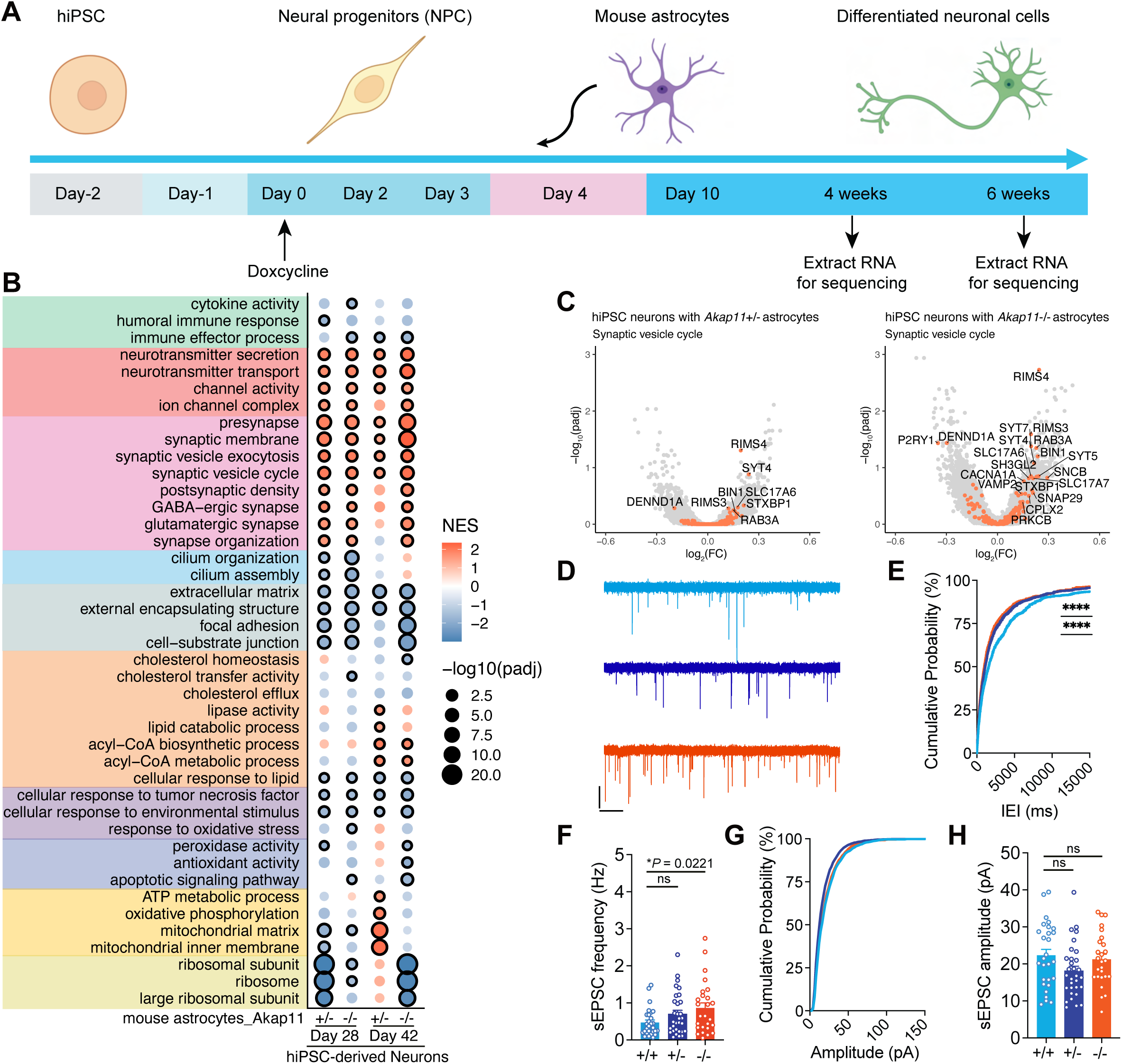
Neuronal activity is increased in hiPSC-derived neurons co-cultured with *Akap11* mutant astrocytes. **(A)** Schematic overview of the co-culture system comprising human iPSC-derived neurons and astrocytes from *Akap11^+/+^*, *Akap11^+/-^*, and *Akap11^-/-^*mice. Neurons were induced by combining Ngn2 expression with forebrain patterning factors. On day 4 of differentiation, neuronal cells were plated onto astrocyte monolayers. RNA was collected at days 28 and 42 of differentiation (DIV28 and DIV42) for transcriptomic analysis. **(B)** Bubble plot of GSEA results highlighting changes in selected molecular pathways. Significant changes (FDR < 0.05) are marked with black outer circles. **(C)** Volcano plot showing significantly increased expression of genes (FDR < 0.05) involved in synaptic vesicle cycle in hiPSC-derived neurons co-cultured with *Akap11* mutant astrocytes at DIV28. Orange dots indicate genes within the synaptic vesicle cycle. **(D)** Representative traces of spontaneous excitatory postsynaptic currents (sEPSCs) at DIV28. (**E and F**) Cumulative probability distribution of sEPSC interevent intervals and average sEPSC frequency recorded at DIV28. Each dot represents an individual culture, N mice: *Akap11*^+/+^ = 5, *Akap11*^+/-^ = 5, *Akap11*^-/-^ = 5). (**G and H**) Cumulative probability distribution of sEPSC amplitudes and average sEPSC amplitude recorded at DIV28. Each dot represents an individual culture, N mice: *Akap11*^+/+^ = 5, *Akap11*^+/-^ = 5, *Akap11*^-/-^ = 5).

GSEA of neurons in the co-cultures showed that *Akap11-*deficient astrocytes, compared to Akap11 wild-type astrocytes, induced significant transcriptional changes in neuronal pathways. Specifically, gene sets related to the extracellular matrix, cilium, immunity, and cellular response to lipid were downregulated, while the acyl-coA metabolic process was upregulated. Notably, the most significantly altered pathways were those related to synapses and neurotransmitter secretion, which were upregulated at both days DIV28 and DIV42 (Figure 6B).

To further characterize the synaptic processes affected in neurons exposed to *Akap11* mutant astrocytes, we performed GO analysis using SynGO-curated terms on the DEGs^92^ (Figure S10C). This analysis revealed significant enrichment of DEGs in pathways related to both the “presynapse” and “postsynapse” (Figure S10C). Further examination identified upregulation of key genes involved in the synaptic vesicle cycle and neurotransmitter release, including *RIMS4*, *RIMS3*, *SYT7*, *SYT4*, *SYT5*, *STXBP1*, and *SNAP29*. Additionally, genes encoding vesicular glutamate transporters, *SLC17A6* (*VGLUT2*) and *SLC17A7* (*VGLUT1*), were upregulated in neurons co-cultured with both *Akap11*^+/-^ and *Akap11*^-/-^ astrocytes compared to *Akap11*^+/+^, with a more pronounced effect observed in the *Akap11*^-/-^ condition (Figure 6C; Table S7), highlighting a potential role for astrocytic Akap11 in regulating neuronal synaptic function.

To determine whether there are functional alterations in synaptic properties, we performed whole-cell patch-clamp recordings on hiPSC-derived neurons at DIV28. Compared to neurons co-cultured with *Akap11*^+/+^ astrocytes, those co-cultured with *Akap11* mutant astrocytes exhibited a gene-dose–dependent increase in the frequency of spontaneous excitatory postsynaptic currents (sEPSCs), while sEPSC amplitude remained unchanged (Figures 6D–6H). To further examine neuronal activity, we conducted calcium imaging on DIV42 hiPSC-derived neuronal co-cultures using the calcium indicator Fluo-4 AM. Calcium signals were minimal in neuron-only and astrocyte-only cultures, and *Akap11* deletion had no effect on calcium activity in astrocyte-only conditions (Figure S11A). Moreover, treatment with tetrodotoxin (TTX), a voltage-gated sodium channel blocker, completely abolished calcium oscillations, confirming that these signals originated from neurons^93^. Consistent with the patch-clamp findings, neurons co-cultured with *Akap11*^-/-^ astrocytes exhibited significantly elevated spontaneous calcium oscillation frequency compared to *Akap11*^+/+^ controls, with *Akap11*^+/-^ astrocytes showing an intermediate increase. In contrast, calcium spike amplitude was unchanged across genotypes (Figures S11B and S11C). To determine whether these functional changes reflected structural differences at synapses, we immunostained for pre- and postsynaptic markers Synapsin and PSD95 at DIV42. Quantification of co-localized Synapsin/PSD95 puncta revealed no significant differences in synapse density across genotypes (Figures S11D and S11E), suggesting that *Akap11*-deficient astrocytes may enhance neuronal activity through an increase in neurotransmitter release probability or intrinsic excitability rather than changes in synapse number or strength. Together, these findings suggest that *Akap11* loss of function in astrocytes enhances neuronal activity compared to *Akap11*^+/+^ controls, through functional changes, rather than structural changes at synapses.

## Discussion

In this study of *Akap11* mutant mice, a potential genetic model of SCZ and BD, we uncovered two key roles for AKAP11 in astrocytes and brain function: 1) *Akap11* deficiency upregulates many genes and proteins involved in lipid metabolism in astrocytes, leading to an accumulation of cholesteryl esters, triacylglycerols, ceramides, and glycerophospholipids, in both astrocytes and brain tissue. 2) Loss of Akap11 in astrocytes increases neuronal activity in co-cultures with hIPSC-derived neurons, which is accompanied by the upregulation of genes involved in synaptic vesicle exocytosis and vesicular glutamate transport. These findings may provide mechanistic insights into AKAP11’s role in SCZ and BD pathophysiology.

Single-nucleus RNA sequencing (snRNA-seq) of the prefrontal cortex from SCZ patients revealed decreased expression of fatty acid and cholesterol biosynthesis/export genes in astrocytes compared to healthy individuals^33^. A similar pattern was observed in *Akap11* mutant mice using snRNA-seq^27^. However, our scRNA-seq data of astrocytes (cultured and isolated from brain) showed upregulation of genes in these pathways (Figures 1B and 2E). This apparent discrepancy could be due to differences in sequencing approach: scRNA-seq captures gene expression changes in the whole cells, while snRNA-seq measures changes within the nucleus. A closer examination of gene-level changes suggests a difference in the affected pathways. In *Akap11* mutant astrocytes, snRNA-seq primarily identifies downregulation of cholesterol synthesis genes, whereas scRNA-seq reveals upregulation of genes involved in cholesterol transport and esterification (Figures 1C and 1D). Some genes, such as *Srebf1*, show consistent upregulation in both snRNA-seq and scRNA-seq datasets (Figures 2C and 2D). While snRNA-seq and scRNA-seq provide both overlapping and distinct biological insights, our lipidomic analysis offers direct evidence that *Akap11* loss-of-function upregulates lipid metabolism, leading to significant intracellular accumulation of cholesteryl esters, triacylglycerols, ceramides, and glycerophospholipids. Intriguingly, AKAP11-mediated lipid regulation is gene-dose dependent, as these lipid elevations were also observed at intermediate level in *Akap11* heterozygous mutant brains (Figures 4C-4H).

Lipid dysregulation has been implicated in both SCZ and BD through other lines of evidence. Genome-wide association studies (GWAS) have identified several possible SCZ risk genes involved in cholesterol metabolism (e.g., *SREBF1*, *SREBF2*, *MSMO1*, *CLU*, *LRP1*, *SOAT2*) and lipid transport or hydrolysis (e.g., *ABCB4*, *ABCB1*, *ABCD2*, *ACADS*, *CPT1C*, *CERS5*, *LIPC*, *ABHD2*)^94–96^. Additionally, analysis of brain gene expression and genetic polymorphisms in SCZ patients has linked *SMPD1* and *SMPD3* to SCZ^97^. Disturbances in fatty acid metabolism have also been observed in *Fads1/2* heterozygous knockout mice; *FADS1/2* is a GWAS-identified BD risk allele^98^. These findings suggest that lipid dysregulation might be a convergent molecular and cellular disturbance shared among various genetic variants associated with SCZ and BD. Beyond genetic evidence, clinical observations further support lipid alterations in SCZ and BD. Patients with SCZ show elevated plasma levels of triglycerides and remnant cholesterol^99^. Increased ceramide levels have also been detected in the white matter of individuals with SCZ and BD^47,48,100^. In *Akap11*^+/-^ mice, we observed an increase in oxidative PUFA products (HODEs, HETEs, HDoHEs) (Figure S6E), which aligns with elevated lipid peroxidation and oxidative stress reported in individuals with SCZ^101–103^.

Interestingly, the aberrant lipid accumulation observed in *Akap11* mutant astrocytes mirrors glial lipid changes in Alzheimer’s disease (AD). In AD, iPSC-derived *APOE4/4* astrocytes and postmortem brains exhibit elevated CE and TG levels alongside matrisome alterations^104,105^. Likewise, AD-associated *APOE4/4* microglia adopt a lipid-droplet–accumulating state marked by ACSL1 upregulation and enhanced TG synthesis^106^. Loss of the microglial receptor TREM2 similarly impairs cholesterol clearance from phagocytic debris, resulting in CE deposition that is reversible by ACAT1 inhibition^107^. Even oligodendrocytes in *APOE4* contexts display aberrant CE deposition associated with compromised myelination^108^. Together, these convergent findings—CE and TG buildup, lipid droplet formation, and disrupted cholesterol transport— define a conserved axis of glial lipid dyshomeostasis in both *Akap11* mutant and AD brains. Notably, APOE4 and TREM2 are among the strongest genetic risk factors for AD, and remarkably, many other AD risk genes (*CLU*, *ABCA7*, *SORL1*, *PICALM*) also function in lipid trafficking or metabolism^109^. Moreover, elevated ceramide levels have been linked to increased AD risk^110–112^ and are relevant to SCZ and BD. Collectively, these findings suggest that altered lipid metabolism may represent a common pathological feature across AD, SCZ, and BD. Although it remains unclear whether these changes are causative or arise as downstream effects of distinct primary mechanisms, they point to potential shared vulnerabilities and new opportunities for therapeutic intervention.

Lipid droplet formation in *Akap11*-deficient astrocytes likely reflects an adaptive buffer against excess lipids and potential lipotoxicity, consistent with upregulation of genes involved in ER stress, oxidative stress, and apoptosis pathways (Figures 1B and 2E). Although astrocytes normally supply cholesterol and other lipids essential for synapse formation and membrane integrity, overloading them can disrupt neuronal function: lipid-laden reactive astrocytes have been shown to promote neuronal hyperactivity and exacerbate epilepsy progression^113^. In our co-cultures, *Akap11* loss likewise increases neuronal activity, although whether this stems from altered lipid delivery, signaling changes, or metabolic stress is still unclear. Future studies should directly test if aberrant lipid trafficking between neurons and astrocytes^89^ drives both lipid droplet formation and neuronal hyperactivity in the context of *Akap11* deficiency.

We observed elevated cAMP levels in *Akap11* mutant mice and astrocytes, along with higher PKA subunit levels and activity (Figure 5), indicating dysregulation of the cAMP/PKA signaling pathway. Interestingly, a similar upregulation of the cAMP pathway has been reported in neurons with a heterozygous loss-of-function mutation in another SCZ risk gene, *SETD1A*^114^. Elevated cAMP-PKA signaling is known to enhance neuronal excitability by modulating ion channels and synaptic plasticity^115,116^. Beyond its role in neuronal excitability, cAMP-PKA signaling is a key regulator of lipid metabolism in various cell types, including astrocytes. Future studies will investigate whether cAMP-PKA activation drives lipid accumulation in *Akap11*-deficient astrocytes.

Technical limitations in this study included the difficulty in quantifying lipid accumulation at spatial resolution *in vivo* and establishing direct causality given the interconnected nature of metabolic pathways. Our co-culture experiments provide insights into how *Akap11*-deficient astrocytes affect neurons, but they may not fully recapitulate the complex interactions in the native brain environment. It remains unclear whether the lipid changes in astrocytes directly cause neuronal hyperactivity or whether other factors are involved. Future studies should generate astrocyte-specific *Akap11* knockout mice to examine how specific metabolic changes contribute to the observed phenotypes, particularly synaptic function and increased neuronal activity. Altogether, our study provides a detailed molecular perspective on AKAP11’s role in astrocytes and the brain, emphasizing its critical involvement in regulating lipid metabolism and cAMP/PKA signaling in astrocytes. These findings enhance our understanding of AKAP11’s biological functions and suggest potential mechanistic links to the pathophysiology of SCZ and BD.

## Acknowledgements

We thank Kris Dickson and members of the Sheng lab for valuable input, and Magdalena Sevilla-Gonzalez for sharing her QC pipeline and input on analysis of metabolomics. We appreciate Fang-Wei Leng’s assistance with visualizing the AKAP11–VAPs interface. We are grateful to Dr. Zhenyu Yue for generously sharing the AKAP11 CRISPR-Cas9 knockout (KO) cell lines derived from HEK293T and HeLa cells.

## Author contributions

X.-M.L. and M.S. conceived and designed the project; X.-M.L. designed, performed, and analyzed most of the experiments in this study; H.P. conducted the initial work and helped with the design and execution of -omics experiments; D.M., K.K., and K.P.M. processed the transcriptome data with guidance from S.K.S. and supervision from J.L; D.G. did the initial work and helped with the design and preparation of cultured astrocyte samples for the -omics experiments; B.J.S conducted the initial characterization of AKAP11; O.S. performed LC-MS/MS for proteomics under the guidance of H.K. and S.A.C.; E.M., C.D., L.D., A.D., and L.I. performed LC-MS for metabolomics under the guidance of C.B.C.; N.D.H. and M.H. performed patch-clamp experiments under the supervision of Z.Y.F.; J.X., M.T., and L.C. provided the hIPSC-derived neurons under the guidance of R.N. and J.Q.P.; Y.-L.Z performed fluorescence Ca^2+^ imaging experiments; W.-C.H. assisted with library preparations for bulk RNA-seq, and J.A. assisted with primary neuron cultures. X.-M.L. and M.S. wrote the manuscript. All authors edited and/or approved the manuscript.

## Methods

### Animals

*Akap11* heterozygous mice (B6.Cg-Akap11[tm1.2Jsco/J]) were obtained from the Jackson Laboratory following cryorecovery (strain #028922). These mice harbor a knockout allele created by deletion of exons 6 and 7 in the mouse *Akap11* gene, as previously described^24^. Prior to deposition, the donor laboratory had backcrossed the mutant strain to C57BL/6J background for six generations. To confirm the genetic background, we conducted SNP analysis using the MiniMuga panel (Transnetyx). Results verified that the *Akap11* mutant mice were >90% C57BL/6J^27^. To establish our colony, founders were crossed to wild-type C57BL/6J mice (Jackson Laboratory, #000664) for one generation. Subsequently, *Akap11* heterozygous mice were intercrossed to produce littermates of three genotypes for experiments: Wild-type (WT, *Akap11*^+/+^), Heterozygous (Het, *Akap11*^+/-^), and Knockout (KO, *Akap11*^-/-^). All animals were housed in AAALAC-approved facilities on a 12-hour light/dark cycle, with food and water available *ad libitum*. All procedures involving *Akap11* mutant mice were approved by the Broad Institute IACUC (Institutional Animal Care and Use Committee) and conducted in accordance with the NIH Guide for the Care and Use of Laboratory Animals. In this study, we used *Akap11* mutant mice and their wild-type littermates at postnatal days 1-4 (P1-P4) and at 1 month of age.

### Cell lines and cell culture

Human induced pluripotent stem cells (iPSCs) were cultured on Geltrex-coated plates (Life Technologies, A1413301) in StemFlex medium (Gibco, A3349401) supplemented with Normocin Antimicrobial Reagent (InvivoGen, Ant-nr-1). Cells were passaged using Accutase (Gibco, A11105).

Human embryonic kidney (HEK293T) cells and human osteosarcoma (U2OS) cells were obtained from the American Type Culture Collection (ATCC). AKAP11 CRISPR-Cas9 knockout (KO) cell lines derived from HEK293T and HeLa cells, along with their corresponding wild-type controls, were generously provided by Dr. Zhenyu Yue (Icahn School of Medicine at Mount Sinai). All cell lines were maintained as monolayer cultures at 37°C in a humidified atmosphere containing 5% CO_2_. Cells were grown in Dulbecco’s Modified Eagle’s Medium (DMEM) supplemented with 10% heat-inactivated fetal bovine serum (FBS) (Life Technologies, 10082147) and 1% penicillin-streptomycin.

### Isolation and culture of mouse astrocytes

Mouse astrocytes were isolated from the cortex and hippocampus of P1-P4 pups as previously described^117^, with modifications. Briefly, the brain was exposed via careful cranial dissection and placed in ice-cold Hank’s Balanced Salt Solution (HBSS). Under a dissecting microscope, the brain was gently stabilized with fine forceps, the two hemispheres were separated, and the cerebellum/brain stem and olfactory bulbs were removed. After meninges removal, the cortices and hippocampi were isolated and cut into small pieces, then transferred to a 15 ml conical tube with HBSS. In a tissue culture hood, HBSS was replaced with 10 ml of pre-warmed plating medium with high glucose DMEM (Gibco, 10569-010) supplemented with 10% heat-inactivated fetal bovine serum (FBS) (Life Technologies, 10082147) and 1% penicillin-streptomycin. The tissue was gently triturated with a 10 ml pipette until the solution was homogeneous. After 1 minute of settling, the cloudy supernatant was centrifuged at 300 x g for 5 minutes. The cell pellet was resuspended in 18 ml of plating medium and transferred to a T75 culture flask. Cultures were incubated at 37°C in a 5% CO_2_ incubator. After 24 hours, the medium was replaced following an HBSS wash. At 10-12 days *in vitro*, microglia were removed by overnight shaking at 37°C, and the medium was replaced. The remaining adherent astrocytes were used for subsequent experiments. In most cases, astrocytes were passaged and cultured in NbActiv4 (BrainBits) media supplemented with 10% heat-inactivated FBS, then shifted to serum-free NbActiv4 media for 3 days. For mass spectrometry analysis, astrocytes were cultured in serum-free DMEM (without phenol red) for 3 days.

### Isolation and culture of rat and mouse hippocampus

Embryonic day 17 (E17) rat or mouse hippocampi were isolated as follows: Pregnant animals were euthanized with CO_2_, and embryos were recovered via cesarean section. Embryos were removed from placental sacs, decapitated at the head/neck junction, and heads were placed in 60 × 15 mm petri dishes containing cold HBSS with 1% penicillin-streptomycin on ice. Under a dissecting microscope, the skull was cut open from caudal to rostral, and the whole brain was removed and placed in a new 60 × 15 mm petri dish with fresh cold HBSS with 1% penicillin-streptomycin. The brain was gently stabilized with fine forceps, the two hemispheres were separated, and the cerebellum/brain stem and olfactory bulbs were removed. After meninges removal, the hippocampus, identified as an opaque, crescent-shaped structure, was excised from each cortical hemisphere using microscissors. Isolated hippocampi were transferred via a trimmed P1000 pipette tip into separate 15 ml tubes containing HBSS with 1% penicillin-streptomycin. The HBSS was then aspirated, and hippocampi were washed thrice with cold PBS. For cell dissociation, hippocampi were incubated in 2 mL of 0.05% trypsin at 37°C for 20 minutes. After trypsin removal and three washes with pre-warmed PBS, 5 ml of pre-warmed NbActiv4 media supplemented with 1% penicillin-streptomycin was added. The tissue was gently pipetted to dissociate neurons, and the resulting suspension was centrifuged at 300 × g for 5 minutes. Following supernatant removal, the cell pellet was gently resuspended in 5 ml of pre-warmed NbActiv4 media with 1% penicillin-streptomycin. Finally, cells were counted and plated on poly-d-lysine coated plates for subsequent analysis.

### Antibodies

The following antibodies were used in this study: a custom polyclonal rabbit anti-AKAP11 antibody was generated by GenScript using the first 284 amino acids of mouse AKAP11 (UniProt: E9Q777) as the antigen sequence (MAAFQPLRSSHLKSKASVRKSFSEDVFRSVKSLLQSEKELCSVSGGECLNQDEHPQLTEVTFLGFNEETDAAHIQDLAAVSLELPDLLNSLHFCSLSENEIICMKDTSKSSNVSSSPLNQSHHSGMLCVMRVSPTLPGLRIDFIFSLLSKYAAGIRHTLDMHAHPQHHLETTDEDDDDTNQSVSSIEDDFVTAFEQLEEEENAKLYNDEINIATLRSRCDAASQTTSGHHLESHDLKVLVSYGSPKSLAKPSPSVNVLGRKEAASVKTSVTTSVSEPWTQRSLY)^27^. Commercial antibodies included: anti-PKA Cα (Cell Signaling Technology, #4782), anti-PKA R1α (Cell Signaling Technology, #5675), anti-VAPA (Proteintech, 15275-1-AP), anti-VAPB (Proteintech, 14477-1-AP), anti-ABCD3/PMP70 (Invitrogen, PA1-650), anti-ATAD3A/3B (Proteintech, 16610-1-AP), anti-PLIN2 (Proteintech, 15294-1-AP), anti-CANX/Calnexin (Abcam, ab22595), anti-LAMP2 (DSHB, H4B4), anti-GFP (Abcam, ab13970), anti-β-actin (Abcam, ab8224), and anti-GAPDH (Cell Signaling Technology, #3683).

### Plasmids and Reagents

The human AKAP11-EGFP construct was generated by cloning the full-length human AKAP11 coding sequence into the pGenLenti_CMV_EGFP vector (GenScript Biotech), incorporating a flexible (GGGGS)₃ linker between AKAP11 and EGFP. EGFP-AKAP11 and mutant variants (EGFP-AKAP11-E354A, EGFP-AKAP11-D357A, EGFP-AKAP11-D360A, and EGFP-AKAP11-E354A/D357A/D360A) were cloned into the pPB_CMV_EGFP vector (VectorBuilder Inc) with a (GGGGS)₃ linker between EGFP and AKAP11.

Transfections were performed using Lipofectamine™ 2000 (ThermoFisher Scientific, Cat# 11668019). The following reagents were used: puromycin (ThermoFisher Scientific, Cat# A1113803), polybrene (Sigma-Aldrich, Cat# H9268), Earle’s Balanced Salt Solution (EBSS; Sigma-Aldrich, Cat# E2888), scramble control siRNA and rat Akap11 siRNA (Integrated DNA Technologies), BODIPY493/503 (Invitrogen™, cat. D3922), IBMX (Sigma-Aldrich, Cat# I5879), GFP-Trap® Agarose (ChromoTek, Cat# gta), and Pierce™ BCA Protein Assay Kit (ThermoFisher Scientific, Cat# 23225).

### Immunohistochemistry

Cells cultured in 96-well plates were washed once with PBS and fixed with 4% EM-grade paraformaldehyde (Electron Microscopy Sciences, Hatfield, PA) for 15 minutes at room temperature (RT). Following fixation, the cells were washed three times with PBS and permeabilized with PBS containing 0.1% Triton X-100 for 10 minutes at RT. After permeabilization, the cells were rinsed three times with PBS and blocked using 100 µL of BlockAid blocking buffer (Thermo Fisher, B10710) for 30 minutes at RT. For antibody staining, the cells were incubated overnight at 4°C with primary antibodies diluted 1:100 or 1:500 in BlockAid buffer. After incubation, the cells were washed three times with PBS (10 minutes per wash) and incubated with secondary antibodies diluted 1:500 in BlockAid buffer for 1 hour at RT. The cells were then washed three additional times with PBS (10 minutes per wash) and stained with DAPI for 10 minutes. Finally, the samples were mounted on slides using ProLong Gold Antifade Reagent (Thermo Fisher). All incubations were performed in the dark to prevent drying and minimize fluorochrome fading. The following primary antibodies were used for immunofluorescence: anti-PKA R1α (Cell Signaling Technology, #5675), anti-p62 (Abcam, ab56416), anti-GFAP (Synaptic Systems, 173004), anti-Synapsin 1/2 (Synaptic Systems, 106006), anti-MAP2 (Synaptic Systems, 188004), and anti-PSD95 (Cell Signaling Technology, #3450S).

### Fluorescence microscopy

Live cell imaging was performed using an LSM900 confocal microscope (Zeiss). Cells were cultured on coverslip-bottom 35 mm dishes (MatTek P35G-1.5-14-C) until reaching approximately 70% confluency. Prior to imaging, cells were stained with 5 μM BODIPY493/503 (Invitrogen™, cat. D3922) or LipidSpot™ 610 (Biotium, cat. 70069) for 30 minutes, followed by 5 μg/ml Hoechst 33342 (Invitrogen™, cat. H3570) for 5 minutes. Images were acquired using a 63× oil-immersion objective (NA 1.4). All image processing and figure preparation were performed using Fiji software (https://fiji.sc/). For fixed samples, cells were cultured in 96-well plates and subjected to standard fixation protocols for immunohistochemistry. Images were acquired on a PerkinElmer Opera Phenix confocal imaging microscope equipped with Cy5, mCherry, GFP, and DAPI filter sets and a 63× 1.4 NA objective lens. Image analysis was conducted using Harmony software. In order to better display the results, we adjusted the ‘scaling’ of the intensity among the panels / mutants in Figure S7H.

### Western blotting

Protein concentrations were determined using Pierce™ BCA Protein Assay Kits (Thermo Fisher Scientific, cat. 23225). Samples were prepared by adding 6x Laemmli buffer (Boston Bioproducts, cat. BP-111R) to a final concentration of 1x and heating at 95°C for 10 minutes. Proteins were separated by SDS-PAGE on 4–12% Novex™ Tris-Glycine gradient gels (Thermo Fisher) and transferred to nitrocellulose membranes using the Trans-Blot Turbo system (Bio-Rad). Membranes were blocked with 5% non-fat milk in Tris-buffered saline containing 0.1% Tween 20 (TBST) and incubated with primary antibodies overnight at 4°C. After three washes in TBST, membranes were incubated with HRP-conjugated secondary antibodies for 1 hours at room temperature. Following three additional 10-minute washes in TBST, protein bands were visualized using the ChemiDoc XRS+ imaging system (Bio-Rad) and quantified with Image Lab Software (Bio-Rad).

### Mass spectrometry analysis of AKAP11 immunoprecipitation

For AKAP11 affinity purification, HEK293T cells stably transfected with AKAP11-EGFP were washed with PBS and resuspended in RIPA buffer (50 mM Tris-HCl, pH 7.5; 150 mM NaCl; 1 mM EDTA; 1% Triton X-100) containing a protease inhibitor cocktail (Roche). Cells were lysed by gentle rotation at 4 °C for 30 minutes, and the lysate was clarified by centrifugation at 13,200 rpm for 15 minutes. The supernatant was incubated with GFP-Trap agarose beads (Chromotek) for 3 hours at 4 °C. After incubation, the beads were washed three times with RIPA buffer and twice with RIPA buffer lacking 1% Triton X-100. Bead-bound proteins were eluted by heating at 95 °C for 10 minutes in Laemmli buffer. For mass spectrometry analysis, the eluted samples were electrophoresed briefly (∼5 minutes) on a 4–20% acrylamide Tris-Glycine gradient gel (Life Technologies). The gel was stained with coomassie blue, and stained bands were excised using a fresh razor blade. To identify total proteins within the excised gel sections, samples were submitted to the Taplin Mass Spectrometry Facility at Harvard Medical School for in-gel tryptic digestion, followed by LC-MS/MS analysis.

### RNA extraction and bulk RNA-seq library preparation

RNA was extracted from cultured primary astrocytes or hiPSC-derived neurons co-cultured with mouse astrocytes using the RNeasy Mini Kit (Qiagen) according to the manufacturer’s protocol. RNA integrity (RIN) was assessed using RNA Pico chips on a 2100 Bioanalyzer Instrument (Agilent), while RNA concentration was determined using a NanoDrop Spectrophotometer. All RNA samples had RIN values greater than 8. Purified RNA samples were stored at −80°C until library preparation for bulk RNA-seq analysis.

Bulk RNA-seq libraries were prepared using the TruSeq Stranded mRNA Kit (Illumina) following the manufacturer’s instructions. For each sample, 200 ng of total RNA was used. The concentration of the resulting cDNA libraries was measured using High Sensitivity DNA chips on a 2100 Bioanalyzer Instrument (Agilent). Normalized libraries (10 nM) were pooled, and sequencing was performed on a NovaSeq S2 platform (Illumina) with 50-base paired-end reads and 8-base index reads.

### Bulk RNA-seq data processing and DE analysis

Raw FASTQ files were aligned to a reference (genome FASTA file and transcriptome GTF were extracted from the CellRanger mm10 reference and used for alignment and quantification) using a Nextflow V1 pipeline^118^ (https://github.com/seanken/BulkIsoform/blob/main/Pipeline). Quantification was performed by Salmon^119^ (version 1.7.0, with arguments −l A --posBias -- seqBias --gcBias --validateMapping), using a Salmon reference built with the salmon index command with genomic decoys included. Quality metrics were obtained from STAR alignment^120^ (version 2.7.10a) and Picard tools CollectRnaSeqMetric command^121^ (version 2.26.7).

Differential expression (DE) analysis was performed comparing the heterozygous or homozygous mutants to wild-type astrocytes for using the Wald test in R package DESeq2^122^ (version 1.34), with batch and sex included as covariates. Salmon output was loaded using R package tximport^123^ (version 1.22). All genes that had at least 10 counts across all samples in each experiment were included in our analysis.

### Purification of mouse astrocytes and single-cell RNA-seq library preparation

One-month-old mice were anesthetized with 3% isoflurane delivered continuously via nose cone, and transcardial perfusions were performed with ice-cold Hank’s Balanced Salt Solution (HBSS, Gibco, 14175-095). The cerebrum was dissected and placed in ice-cold HBSS. Astrocytes were isolated and enriched using the Adult Brain Dissociation Kit, mouse and rat (Miltenyi, 130-107-677) and the Anti-ACSA-2 MicroBead Kit, mouse (Miltenyi, 130-097-679) following the manufacturer’s instructions with modifications. Briefly, brain tissue was enzymatically digested in a C tube (Miltenyi, 130-093-237) with Enzyme Mix 1 and 2, then processed using a gentleMACS Octo Dissociator with Heaters (program 37C_ABDK_02 for 20-100 mg tissue, or 37C_ABDK_01 for >100 mg). After centrifugation (300 × g, 2 min, 4°C), the sample was resuspended in 1 mL cold D-PBS (Gibco, 14287-080) and filtered through a 70 μm strainer (Miltenyi MACS SmartStrainers, 130-110-916). The C-tube was rinsed with 10 mL cold D-PBS and applied to the filter. Following centrifugation (300 × g, 10 min, 4°C), the pellet was resuspended in 3100 μL cold D-PBS with 900 μL cold Debris Removal Solution (Miltenyi, 130-109-398), overlaid with 4 mL cold D-PBS, and centrifuged (3000 × g, 10 min, 4°C). The top two phases were removed, and the remaining solution was diluted to 7.5 mL with cold D-PBS, inverted gently, and centrifuged (1000 × g, 10 min, 4°C). The pellet was resuspended in PB buffer (0.5% BSA, 0.2 U/μL RNase inhibitor, 1× PBS) and incubated with FcR blocker (10 min, 4°C) followed by ACSA-2 microbeads (15 min, 4°C) in the dark. For each hemisphere, 160 μL cold PB buffer, 20 μL FcR blocker, and 20 μL ACSA-2 beads were used. The cell-bead mixture was diluted to 500 μL with cold PB buffer and applied to a prepared MS column. After three 500 μL PB buffer washes, cells were eluted with 1 mL cold PB buffer. Isolated cells were centrifuged (300 × g, 5 min, 4°C), resuspended in 100 μL cold PB buffer, and counted using a Cellometer K2 with AOPI dye (15 μL sample + 15 μL dye, 20 μL loaded onto slide). Cell viability and total cell count were recorded.

For single-cell RNA-seq library preparation, 16,500 cells per sample were loaded into the 10x Chromium V3.1 system (10x Genomics) and processed according to the manufacturer’s protocol. The resulting 10 nM normalized library was pooled, and sequencing was performed on a NovaSeq S2 (Illumina) for two rounds with 28 and 75 bases for reads 1 and 2, respectively, and 10 bases each for index reads 1 and 2.

### ScRNA-seq data processing and analysis

#### ScRNA-seq data analysis pipeline

ScRNA-seq binary cell call (BCL) files were demultiplexed to FASTQ reads for *Akap11^+/+^*, *Akap11^+/-^*, and *Akap11^-/-^*mouse samples using the mkfastq function from Cell Ranger v7.2.0 with the default parameters^124^. Reads were aligned to [mm10] reference transcriptome. Gene by cell UMI count matrices were obtained using the count function of the Cell Ranger. R version 4.2.2, and Seurat R package version 4.3.0 was used for the downstream analyses. For the initial quality filtering, cells expressing more than 500 genes, but less than 20000 were kept. Cells with greater than 10% mitochondrial gene expression were removed. UMI counts for each cell were normalized by the total counts for that cell, multiplied by a scaling factor of 10000 and then natural log transformed. Variable features (genes) were identified using the variance stabilizing transformation method with default parameters. The Seurat’s ScaleData function was used to regress out variation due to differences in total UMIs per cell and percent mitochondrial gene expression. Doublets were identified and removed with Scrublet with doublet threshold set to 0.5. Principal component analysis (PCA) was performed on the scaled data for the variable genes. The top 30 PCs were used to construct shared nearest neighbor (SNN) graph by calculating the neighborhood overlap (Jaccard index) and reduce dimensionality using the uniform manifold approximation and projection (UMAP). Cells were then clustered on the SNN graph using the Seurat’s FindCluster function with resolution parameter set to 0.3.

#### Differentially expressed genes

To identify differentially expressed (DE) genes in each cell cluster and condition, we employed a pseudo-bulk likelihood ratio test approach using the DESeq2 R package (version 1.38.3). Lowly expressed genes (<10 supporting reads or <0.1 normalized expression) were identified within each comparison and excluded from the DE test. Computed p-values were adjusted using the Benjamini-Hochberg method.

#### scRNA-seq cluster annotations

Cell type for each cluster was initially assigned using MapMyCells^37^, a web application from Allen Brain Institute designed to label cell types using a reference dataset. 10x Genomics whole mouse brain dataset (CCN20230722) was used as the reference, and hierarchical mapping method was used. Query dataset was down-sampled to 500 cells per cluster. The initial cell type labels were then verified using the cluster-specific DE genes.

#### scRNA-seq astrocyte subset data processing

Identified astrocytes were isolated and processed through the scRNA-seq data analysis pipeline described above, except for PCA. Instead, the dimensionality reduction was performed using scGBM^125^, which tries to better capture relevant biology in its low-dimensional latent factors by directly modeling the count. The top 20 latent factors were used for the downstream analyses in the pipeline. One low quality cluster (<2000 mean UMI count and <1000 number of genes) and another cluster with higher immediate early genes (IEG) (Supplementary Table 6) expression were identified and excluded from further annotation of brain regions. IEG enriched cluster was identified using Seurat’s AddModuleScore function. The remaining astrocytes were processed and labeled with brain regions using the same annotation method described in the previous section.

### Liquid chromatography tandem mass spectrometry (LC-MS/MS) proteomics

#### Protein digestion

Cultured astrocytes were lysed in a buffer containing 2% SDS containing protease and phosphatase inhibitors. Lysates were then reduced with 5 mM dithiothreitol (DTT, Sigma-Aldrich) at 37°C (1000 rpm, thermomixer) for 1 hour, and alkylated with 10 mM iodoacetamide (IAA, Sigma-Aldrich) at 25°C (1000 rpm, thermomixer) for 45 minutes in the dark. Samples were subsequently acidified with phosphoric acid (final concentration 1.2%). Acidified samples were mixed 1:6 with chilled S-trap buffer (90% methanol, 100 mM triethylammonium bicarbonate (TEAB)) and loaded onto midi S-trap columns (Protifi, Fairport, NY). The S-trap protocol was then followed: samples were centrifuged for 1 minute at 4000 rcf, washed three times with 3 mL of S-trap buffer (each time centrifuged at 4000 rcf), and transferred to clean 15 mL tubes. Digestion was performed by adding 350 μL digestion buffer containing LysC endopeptidase (enzyme:substrate ratio 1:40, 1mAU/μL, Wako Chemicals, Richmond, Virginia) and sequencing-grade modified trypsin (enzyme:substrate ratio 1:40, Promega). Spin once more, the collected digestion buffer flow-through was pipetted back onto the column, and digestion proceeded overnight at room temperature (approximately 16 hours). Columns were then eluted sequentially with 500 µL each of 50 mM TEAB, 0.1% formic acid, and 500 µL 0.1% formic acid 50% acetonitrile, with 1-minute centrifugations at 4000 rcf between each elution step. Eluates were combined, frozen, dried using a vacuum concentrator, resuspended in 3 mL of 0.1% formic acid with 3% acetonitrile, and desalted using Sep-Pak C18 SPE cartridges (Waters, Milford, MA). Peptides were dried using a vacuum concentrator, and their concentrations were determined by BCA assay. Subsequently, 250 µg aliquots were prepared for tandem mass tag (TMT) labeling.

#### Tandem mass tag (TMT) labeling of peptides

Most replicates within *Akap11*^+/+^, *Akap11*^+/-^ and *Akap11*^-/-^ groups were biological replicates belonging to 2 different cohorts. To increase the total number of replicates for a few samples, technical replicates belonging to the same digested sample were included. Aliquots containing 250 μg of digested peptides were resuspended in 50 mM HEPES (pH 8.5) at a final concentration of 5 μg/μL for TMT labeling. Experimental samples were randomized and assigned across the 16 TMT labeling channels (126C–134N). Each sample was combined with 250 μg of the respective TMT reagent and incubated for 1 hour at 25°C (1000 rpm, thermomixer). Samples were then diluted to 2 μg/μL in 50 mM HEPES, a small aliquot was removed from each sample for labeling efficiency and mixing tests, and the remaining sample was snap-frozen in liquid nitrogen and stored at −80°C. Following quality-control tests, labeling reactions were quenched using 5% hydroxylamine, and samples within each multiplex set were pooled. Pooled samples were dried in a vacuum concentrator and desalted using Sep-Pak C18 SPE cartridges (Waters).

#### Offline bRP fractionation

Each combined multiplex sample was fractionated by high pH reversed phase separation using a 3.5 μm Agilent Zorbax 300 Extend-C18 column (4.6 mm ID x 250 mm length). Samples were resuspended in mobile phase A (5 mM ammonium formate, pH 10, in 2% acetonitrile), centrifuged to remove debris, loaded onto the column and eluted off the column for 96 minutes at a flow rate of 1 ml/minute using mobile phase B (5 mM ammonium formate, pH 10, in 90% acetonitrile) with the following gradient (time(min): %B): 0:0; 7:0; 13:16; 73:40; 77:44; 82:60; 96:60. A total of 96 fractions were collected and concatenated down to 24 fractions by combining non-sequential fractions. Five percent was removed for global proteome analysis and the remaining 95% was further concatenated to 12 fractions for phosphopeptide enrichment. Fractions were snap-frozen in liquid nitrogen, dried using a vacuum concentrator, and stored at −80°C until phosphopeptide enrichment.

#### Phosphopeptide enrichment

Phosphopeptide enrichment was performed using Agilent AssayMap BRAVO Sample Prep platform through immobilized metal affinity chromatography (IMAC) with AssayMap Fe3+-NTA cartridges (Agilent, Santa Clara, California). Peptide fractions were resuspended in 200 μL 80% ACN + 0.1% TFA. After thorough reconstitution, samples were centrifuged for 5 minutes at 6,000 rcf to pellet insoluble material, and 190 μL was transferred to the plate for BRAVO IMAC enrichment. BRAVO instrument was run with the a priming/syringe wash buffer of 50% acetonitrile 0.1% TFA, elution buffer of 1% ammonium hydroxide, equilibration/cartridge wash buffer of 80% acetonitrile 0.1% TFA, and the following settings: 250 μL 3x initial syringe washes, 100 μL at 300 μL/min 1x cartridge priming, 50 μL at 10 μL/min 1x cartridge equilibration, 350 μg sample amount in 190 μL at 5 μL/min 3x, flow through collection checked, 25 μL 1x cup wash, 75 μL at 10 μL/min 3x internal cartridge wash, flow through collection checked, 50 μL 1x stringent syringe wash, 20 μL at 5 μL/min 1x sample elution, 0 μL eluate discard, 2.5 μL 100% formic acid existing collection volume, and 250 μL 3x final syringe wash. The flow through plate was frozen and stored at −80°C. Eluted peptides were transferred to MS vials, dried using a vacuum concentrator, and reconstituted in 9 μl of 0.1% FA and 3% ACN.

#### LC-MS/MS analysis of samples

Online separation was conducted with a Vanquish Neo UHPLC system (Thermo Fisher Scientific). An in-house packed 26 cm x 75 μm internal diameter C18 silica picofrit capillary column (1.9 mm ReproSil-Pur C18-AQ beads, Dr. Maisch GmbH, r119.AQ; Picofrit 10 μm tip opening, New Objective, PF360-75-10-N-5) heated at 50°C using a column heater sleeve (Phoenix-ST) was used for chromatographic separation. For global proteome, 1 μg was loaded on-column. For phosphoproteome, 50% of each fraction was injected in a 4-μl volume. Flow rate was 200 nL/min, and mobile phases were composed of 0.1% formic acid / water for solvent A and 0.1% formic acid / Acetonitrile for solvent B. The 110-minute liquid chromatography (LC) method used for global proteome and IMAC phosphoproteome analysis consisted of a 10-minute column-equilibration procedure; a 20-minute sample-loading procedure; and the following gradient profile: (time (minutes):%B) 0:1.8; 1:5.4; 85:27; 94:54; 95;81; 100:81; 101:45; 110:45. All the samples were analyzed on an Orbitrap Exploris 480 mass spectrometer (Thermo Fisher Scientific). Data-dependent MS/MS acquisition was performed using the following relevant parameters: positive ion mode at a spray voltage of 1.8 kV, MS1 resolution of 60,000 with a normalized AGC target of 300% and maximum inject time of 10ms, a mass range from 350 to 1800 m/z, 20 most abundant ions selected with MS2 resolution of 45,000, MS2 AGC target of 30% and MS2 maximum injection time of 105 ms, isolation window of 0.7 m/z, instrument specific HCD collision energy of 32 or 34%, dynamic exclusion for 15 seconds. Precursor fit threshold was set to 50% with a fit window of 1.2.

#### Raw MS/MS data processing

The data was analyzed using Spectrum Mill MS Proteomics Software (Broad Institute) with a mouse database from Uniprot.org downloaded on 04/07/2021 containing 55734 entries. Search parameters included: ESI Q Exactive HCD v4 35 scoring parent and fragment mass tolerance of 20 ppm, 40% minimum matched peak intensity, trypsin allow P enzyme specificity with up to four missed cleavages and calculate reversed database scores enabled. Fixed modifications were carbamidomethylation at cysteine. TMT labeling was required at lysine, but peptide N termini could be labeled or unlabeled. Allowed variable modifications were protein N-terminal acetylation, oxidized methionine pyroglutamic acid, and pyro carbamidomethyl cysteine, as well as phosphorylated serine, threonine, and tyrosine for the phosphoproteome dataset. Protein quantification was achieved by taking the ratio of TMT reporter ions for each sample over the TMT reporter ion for the median of all channels. TMT16 and TMT18 reporter ion intensities were corrected for isotopic impurities in the Spectrum Mill protein/peptide summary module using the afRICA correction method which implements determinant calculations according to Cramer’s Rule and correction factors obtained from the reagent manufacturer’s certificate of analysis (https://www.thermofisher.com/order/catalog/product/90406) for lot numbers XC342533 and XI346567 / XJ346678 for lysate and secretome experiments, respectively. Log2 ratios for proteins and phosphosites were exported from Spectrum Mill and median-MAD normalization was applied to all datasets prior to statistical analysis.

#### Statistical analysis

All batch correction and linear modeling was performed using the limma package in R. Batch correction for cohort was performed on the log2-transformed ratio values after median MAD normalization and prior to linear modeling using the removeBatchEffect function. Genotype was included in the design matrix for batch correction to retain differences due to genotype. After batch correction, features with >30% missing values were removed prior to linear modeling. The linear model contained a fixed effect for genotype and a random effect for the technical replicates which was incorporated using the duplicateCorrelation function. Empirical Bayes shrinkage was applied prior to calculating fold changes and p-values for each pairwise contrast. Protein NNT was excluded from the downstream analysis, as it represents a background mutation in the mouse strain that coincidentally segregated with the Akap11 genotype in this experiment.

### Lipidomic and metabolomic analysis of brain tissue and astrocyte cultures

Metabolomics on brain tissue and astrocyte cultures was performed using a platform that relies on a combination of four non-targeted liquid chromatography mass spectrometry (LC-MS) methods which measure both polar and non-polar metabolites^126^. For cultured astrocytes (grown in 6-well plates), extracts were generated from approximately 750,000 cells per well using either 0.8 mL methanol (polar extraction) or 1 mL isopropanol (non-polar extraction). Brain tissues were homogenized using a Qiagen TissueLyzer II in water at 1:4 (w:v) (TissueLyser II; Qiagen Inc.; Valencia, ca) and the aqueous homogenate was subjected to protein precipitation. Samples were prepared for each method using extraction procedures that are matched for use with the chromatography conditions. Data were acquired using LC-MS systems comprised of Nexera X2 U-HPLC systems (Shimadzu Scientific Instruments) coupled to Q Exactive/Exactive Plus orbitrap mass spectrometers (Thermo Fisher Scientific). The method details are summarized below.

#### LC-MS Method 1: HILIC-pos (positive ion mode MS analyses of polar metabolites)

LC-MS samples were prepared from media (10 μL) by protein precipitation with the addition of nine volumes of 74.9:24.9:0.2 v/v/v acetonitrile/methanol/formic acid containing stable isotope-labeled internal standards (valine-d8, Isotec; and phenylalanine-d8, Cambridge Isotope Laboratories). The samples were centrifuged (10 min, 9,000 × g, 4°C), and the supernatants injected directly onto a 150 × 2-mm Atlantis HILIC column (Waters). The column was eluted isocratically at a flow rate of 250 μL/min with 5% mobile phase A (10 mM ammonium formate and 0.1% formic acid in water) for 1 min followed by a linear gradient to 40% mobile phase B (acetonitrile with 0.1% formic acid) over 10 min. MS analyses were carried out using electrospray ionization in the positive ion mode using full scan analysis over m/z 70–800 at 70,000 resolution and 3-Hz data acquisition rate. Additional MS settings are: ion spray voltage, 3.5 kV; capillary temperature, 350°C; probe heater temperature, 300°C; sheath gas, 40; auxiliary gas, 15; and S-lens RF level 40.

#### LC-MS Method 2: HILIC-neg (negative ion mode MS analysis of polar metabolites)

LC-MS samples were prepared from media (30 μL) by protein precipitation with the addition of four volumes of 80% methanol containing inosine-15N4, thymine-d4 and glycocholate-d4 internal standards (Cambridge Isotope Laboratories). The samples were centrifuged (10 min, 9,000g, 4°C) and the supernatants were injected directly onto a 150 × 2.0-mm Luna NH2 column (Phenomenex). The column was eluted at a flow rate of 400 μL/min with initial conditions of 10% mobile phase A (20 mM ammonium acetate and 20 mM ammonium hydroxide in water) and 90% mobile phase B (10 mM ammonium hydroxide in 75:25 v/v acetonitrile/methanol) followed by a 10-min linear gradient to 100% mobile phase A. MS analyses were carried out using electrospray ionization in the negative ion mode using full scan analysis over m/z 60–750 at 70,000 resolution and 3 Hz data acquisition rate. Additional MS settings are: ion spray voltage, −3.0 kV; capillary temperature, 350°C; probe heater temperature, 325°C; sheath gas, 55; auxiliary gas, 10; and S-lens RF level 40.

#### LC-MS Method 3: C18-neg (negative ion mode analysis of metabolites of intermediate polarity; for example, bile acids and free fatty acids)

Media homogenates (30 μL) were extracted using 90 μL methanol containing PGE2-d4 as an internal standard (Cayman Chemical Co.) and centrifuged (10 min, 9,000g, 4°C). The supernatants (10 μL) were injected onto a Acquity HSS T3 C18 2.1 x 150mm (1.8um; Waters). The column was eluted isocratically at a flow rate of 450 μL/min with 20% mobile phase A (0.01% formic acid in water) for 3 min followed by a linear gradient to 100% mobile phase B (0.01% acetic acid in acetonitrile) over 12 min. MS analyses were carried out using electrospray ionization in the negative ion mode using full scan analysis over m/z 70–850 at 70,000 resolution and 3 Hz data acquisition rate. Additional MS settings are: ion spray voltage, −3.5 kV; capillary temperature, 320°C; probe heater temperature, 300°C; sheath gas, 45; auxiliary gas, 10; and S-lens RF level 60.

#### LC-MS Method 4: C8-pos

Lipids (polar and nonpolar) were extracted from media (10 μL) using 190 μL isopropanol containing 1-dodecanoyl-2-tridecanoyl-sn-glycero-3-phosphocholine as an internal standard (Avanti Polar Lipids; Alabaster, AL). After centrifugation (10 min, 9,000g, ambient temperature), supernatants (10 μL) were injected directly onto a 100 × 2.1-mm ACQUITY BEH C8 column (1.7 μm; Waters). The column was eluted at a flow rate of 450 μL/min isocratically for 1 min at 80% mobile phase A (95:5:0.1 v/v/vl 10 mM ammonium acetate/methanol/acetic acid), followed by a linear gradient to 80% mobile phase B (99.9:0.1 v/v methanol/acetic acid) over 2 min, a linear gradient to 100% mobile phase B over 7 min, and then 3 min at 100% mobile phase B. MS analyses were carried out using electrospray ionization in the positive ion mode using full scan analysis over m/z 200–1,100 at 70,000 resolution and 3 Hz data acquisition rate. Additional MS settings are: ion spray voltage, 3.0 kV; capillary temperature, 300°C; probe heater temperature, 300°C; sheath gas, 50; auxiliary gas, 15; and S-lens RF level 60.

#### Metabolomics data processing

Nontargeted data were processed using Progenesis QI software (v 2.0, Nonlinear Dynamics) to detect and de-isotope peaks, perform chromatographic retention time alignment, and integrate peak areas. Peaks of unknown identity were tracked by method, m/z and retention time. Identification of the resulting nontargeted metabolite LC-MS peaks was conducted by: i) matching measured retention times and masses to mixtures of reference metabolites analyzed in each batch; and ii) matching an internal database of compounds that have been characterized using the Broad Institute methods. Temporal drift was monitored and normalized with the intensities of peaks measured in the pooled reference samples.

To identify differentially expressed (DE) lipids and metabolites, data from each method were merged and subjected to a quality control (QC) pipeline. This pipeline included: removal of internal standards and within-method duplicates, exclusion of metabolites with >25% missing values, imputation of missing values using half of the minimum value, windsorization to ±5 standard deviations, and log_2_ transformation. For the Shiny GATOM integrative analysis shown in Figure S5A, inverse normal transformation was applied instead. Differential analysis was performed using the limma model, with sex as a covariate.

### Neuronal differentiation and co-culture with mouse astrocytes

Human induced pluripotent stem cells (hiPSCs) from the WTC11 line (for day 28) or PGP1 line (for day 42) were differentiated into cortical glutamatergic neurons following a previously described protocol with modifications^90^. hiPSCs were seeded at 80,000 cells/cm^2^ in mTeSR medium supplemented with 10 µM Y-27632 (Stemgent, 04-0012), 2 µg/ml doxycycline hyclate and 0.2% Normocin. On day 2, the medium was replaced with Neural Induction Medium (NIM) consisting of DMEM/F12 supplemented with 1% N2, 1% GlutaMAX, 1.5% D-(+)-glucose, and 0.2% Normocin. NIM was further supplemented with 10 µM SB431542 (Tocris, 1614), 2 µM XAV939 (Stemgent, 04-00046), and 100 nM LDN-193189 (Stemgent, 04-0074). Doxycycline hyclate (2 μg/mL) was maintained throughout the differentiation process unless otherwise noted. On day 3, cultures were maintained in the same medium with the addition of cytosine β-D-arabinofuranoside (AraC). On day 4, the medium was switched to Neurobasal medium (Gibco) supplemented with 2% B27 (Gibco), 0.5% MEM Non-Essential Amino Acids, 1.5% D-(+)-glucose, 1% GlutaMAX, and 0.2% Normocin. This medium was further supplemented with 10 ng/ml each of brain-derived neurotrophic factor (BDNF), ciliary neurotrophic factor (CNTF), and glial cell-derived neurotrophic factor (GDNF) (all from R&D Systems; 248-BD/CF, 257-NT/CF, and 212-GD/CF, respectively).

For co-culture experiments, day 4 neuronal progenitors were dissociated using accutase and seeded at 80,000 cells/cm^2^ onto geltrex-coated plates containing mouse primary cortical glial cells. Primary glial cells were prepared from the cortex and hippocampus of postnatal days 1-4 (P1-P4) mouse pups as previously described.

### Bulk RNA-seq data processing and analysis from co-cultures hiPSC-derived neurons and mouse astrocytes

#### Bulk RNA-Seq data alignment

Raw sequencing reads (FASTQ files) from mixed-species co-cultures of hiPSC-derived neurons and mouse astrocytes were aligned to a custom combined human/mouse reference. This reference was constructed using Salmon^119^ (v1.4.0) with the ‘salmon index’ command, using concatenated human (GRCh38.p14) and mouse (GRCm39) genome FASTA and GTF files from GENCODE^127^. Alignment and quantification were performed using Salmon with the following arguments: −l A --posBias --seqBias --gcBias --validateMapping. To assess the accuracy of species-specific read mapping, control single-species samples (hiPSCs and mouse astrocytes) were aligned to the same combined reference used for mixed-species co-cultures. For the control samples, reads mapped to the correct species with high accuracy: approximately 99.6% for human-only hiPSC samples and 96% for astrocyte-only mouse samples (Figure S10). One control single-species astrocyte sample (H1HT) was excluded from analysis due to hiPSC contamination. In all samples, reads mapping to multiple genomic locations were removed from the analysis. The custom reference construction and read alignment were performed using a NextFlow V1 pipeline (https://github.com/seanken/BulkIsoform/blob/main/Pipeline)^118^.

#### Differential Expression (DE) analysis

DEGs were identified across *Akap11* genotypes in the mixed-species co-cultured samples as follows. Confidently assigned reads were imported using the R package tximport (v1.28.0)^123^. Reads for each genetic replicate were combined (4 replicates per mouse, 18 mice total), and reads aligned to human and mouse genes were separated into different data frames. Principal Component Analysis (PCA) plots showing separation of single species cultured hiPSC neurons (controls) and mixed species co-cultured hiPSC neurons were generated using the R packages ggplot2 (v3.5.1) and cowplot (v1.1.3) (Figure S10). Lowly expressed genes were filtered out using the *filterbyexpr()* function from the edgeR package (v3.42.4). DESeq2 (v1.40.2)^122^ was used to identify differentially expressed genes in hiPSC-derived neurons co-cultured with *Akap11* mutant mouse astrocytes (*Akap11*^-/-^ vs *Akap11*^+/+^, and *Akap11*^+/-^ vs *Akap11*^+/+^). Sex was included as a covariate to account for astrocytes derived from both male and female mice. Additionally, the log_10_ value of the percentage of reads confidently assigned to each species per sample was included as a covariate to account for differences in species-specific read proportions.

Log_2_ fold change shrinkage was performed on DE results using DESeq2’s *lfcShrink()* tool with ‘normal’ shrinkage estimation^122^. This analysis was conducted separately for human and mouse gene counts, with similar contrasts run for astrocytes in mixed-culture samples. DEGs were defined as those with a FDR < 0.05.

### Gene set enrichment analysis

GSEA was performed with the R Bioconductor package fgsea^123^ (version 1.26.0) using C5 ontology gene sets obtained from the MsigDB. Genes were ranked using DESeq2’s Wald statistic (stat), and the analysis was conducted separately for each differential expression contrast. The Gene Ontology (GO) database was used as the source of gene sets, with minimum and maximum gene set sizes set to 15 and 500, respectively, as recommended by the fgsea package. Gene sets were obtained using the R package AnnotationDbi (v1.56.2). GSEA results across comparisons were compiled and visualized using the R packages dplyr (v1.1.3) and ggplot2 (v3.5.1). Additionally, synapse-specific gene ontology (synGO) analysis was performed on differentially expressed genes using the synGO gene set database (v1.1)^92^. Significantly enriched synapse-related gene sets (defined by FDR < 0.05) were visualized using the online synGO plotter tool (https://www.syngoportal.org/plotter).

### LANCE ultra cAMP assay

The cAMP assay was performed using the LANCE Ultra cAMP Assay Kit (PerkinElmer, catalog TRF0262) following the manufacturer’s protocol. Brain tissues were homogenized in RIPA buffer (50 mM Tris-HCl, pH 7.5; 150 mM NaCl; 1 mM EDTA; 1% Triton X-100) containing a protease inhibitor cocktail, using a Qiagen TissueLyzer II, and subsequently diluted in stimulation buffer (1× HBSS, 5 mM HEPES, 0.5 mM IBMX, 0.1% BSA, pH 7.4). Cell lysates were similarly prepared in RIPA buffer with protease inhibitors and diluted in the same stimulation buffer. Protein concentrations were determined using Pierce™ BCA Protein Assay Kits.

For each assay reaction, 5 μL of cell lysate was combined with 5 μL of stimulation buffer, 5 μL of 4× Eu-cAMP tracer working solution, and 5 μL of 4× ULight-anti-cAMP working solution. The mixture was incubated at room temperature for 1 hour, sealed with TopSeal-A film, and read on a TR-FRET microplate reader. All assays were performed in triplicate. cAMP levels were normalized to total protein concentration.

### Whole-cell electrophysiology

Cover slips containing hiPSC neurons and mouse astrocyte co-cultures at 4 or 6 weeks *in vitro* were transferred to a recording chamber and perfused at 1-2 mL/min with an extracellular solution containing (in mM): 119 NaCl, 2.3 KCl, 2 CaCl_2_, 1 MgCl_2_, 15 HEPES, 5 glucose, cresol red (0.25 mg/L), and D-serine (10 μM). Extracellular solution pH was adjusted to 7.2-7.4 with NaOH and osmolality adjusted to ∼325 mOsm with sucrose. Picrotoxin (50 μM) was added to the solution to block fast GABA_A_ receptor-mediated inhibitory neurotransmission. Neurons were visually identified under a 40X objective using differential interference contrast (DIC) microscopy. Whole-cell patch clamp recordings of hiPSC neurons were made at room temperature using glass capillaries (KG33, King Precision Glass) with 3-6 MΩ tip resistance filled with an intracellular solution containing (in mM): 110 Cs-Gluc, 4 NaCl, 15 KCl, 5 TEA-Cl, 20 HEPES, 0.2 EGTA, 5 lidocaine N-ethyl bromide, 4 ATP magnesium salt, and 0.3 GTP sodium salt. Intracellular solution pH was adjusted to 7.2-7.4 with CsOH and osmolality adjusted to 298-300 mOsm with ∼ 10 mM K_2_SO_4_. Neurons were voltage-clamped at −70 mV using a multiclamp 700B amplifier (Molecular Devices) and a 10 mV square pulse was used to monitor membrane properties online. Intracellular solution was allowed to dialyze for ∼5 min to allow sufficient block of potassium channels and membrane properties to stabilize. Spontaneous excitatory post synaptic currents (sEPSCs) were then recorded for 3 minutes. Neurons in which the access resistance (R_a_ typically between 8-12 MΩ) was greater than 25 MΩ or changed by >20% during the recording were excluded from analysis. All sEPSC recordings were sampled at 10 KHz and low pass filtered at 2 KHz using a Digidata 1440A digitizer (Molecular Devices), stored offline, and analyzed using ClampFit software version 10.6 (pClamp Software suite, Molecular Devices). The average sEPSC frequency and amplitude was calculated and plotted using GraphPad version 10 (Prism) and statistical outliers were excluded from analysis using ROUT’s test.

### Fluorescence Ca^2+^ imaging with Fluo-4 AM

Human iPSC-derived neurons (6k/well) and mouse astrocytes (3.5k/well) were co-cultured in Poly-D-Lysine-coated 384-well plates (Corning BioCoat) for 6 weeks. Before imaging, the culture medium was removed, and cells were washed with 25 μL of buffer A (HBSS/20 mM HEPES, pH 7.3). The co-culture was then loaded with 25 μL of 2 μM Fluo-4 AM (Thermo Fisher Scientific, Catalog No. F14201) in buffer A and incubated for 2 hours in the dark at room temperature. Cells were gently washed twice with 25 μL of buffer A, and 25 μL of buffer A was then added to each well. Plates were placed in the FLIPR® Penta (Molecular Devices), and calcium images of all 384 wells were captured simultaneously every second for a total of 4 minutes at 37°C, with excitation at 470–495 nm and emission at 515–575 nm.

## Data and software availability

The original mass spectra, spectral library, and the protein sequence databases used for searches have been deposited in the public proteomics repository MassIVE (http://massive.ucsd.edu) and are accessible at ftp://MSV000097493@massive.ucsd.edu. These datasets will be made public upon acceptance of the manuscript. Raw FASTQ files from bulk and single-cell RNA sequencing experiments will be deposited at NCBI GEO. Original metabolomics raw data will be uploaded to the Metabolomics Workbench. All datasets will become publicly available upon manuscript acceptance. Software used for data analysis is publicly accessible.

---

**Figure S1.**
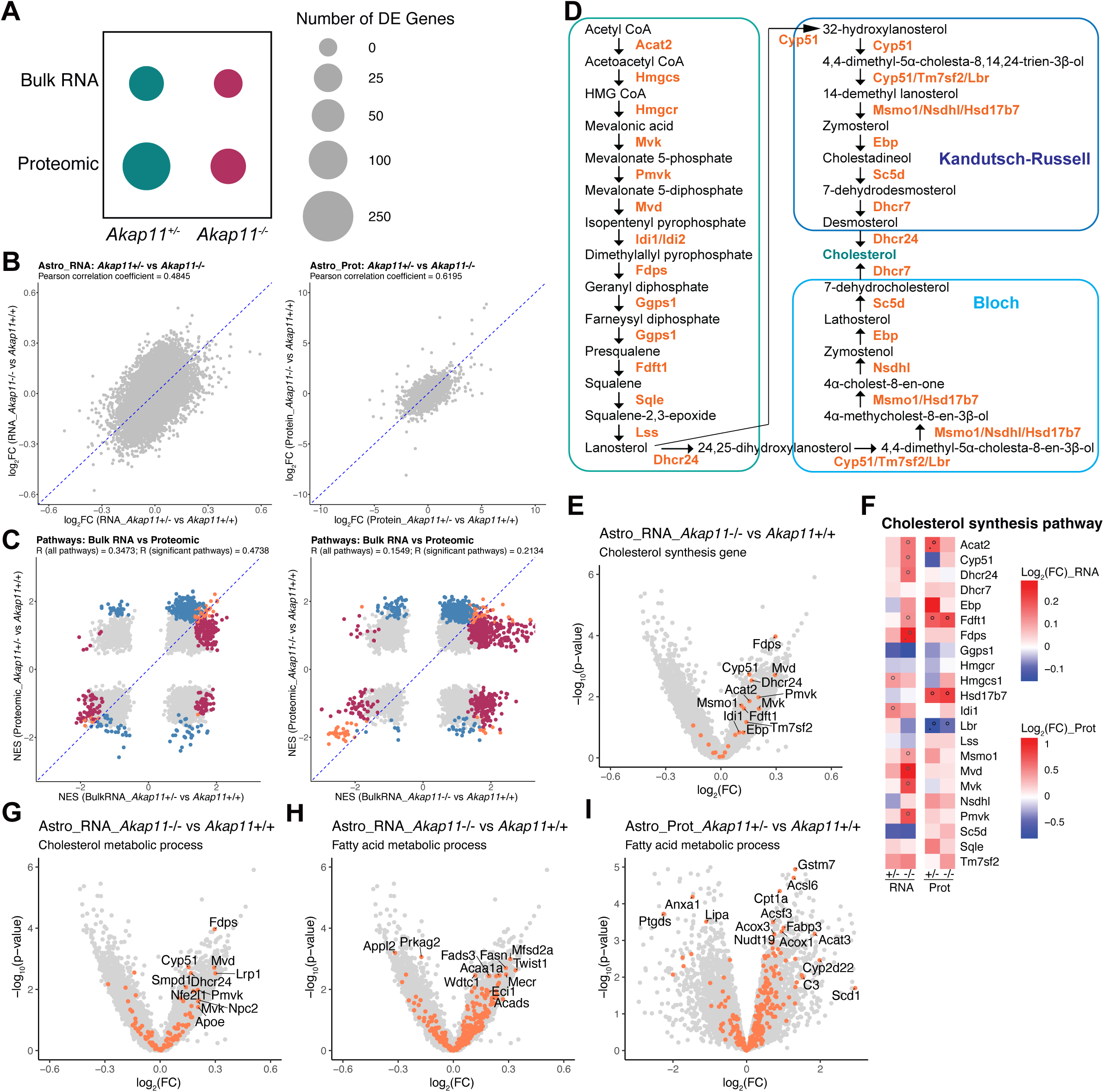
Integrated transcriptomic and proteomic analysis reveals altered lipid metabolic pathways in *Akap11* mutant astrocytes. (A) Number of differentially expressed genes (DEGs) and differentially expressed proteins (DEPs) in *Akap11*^+/-^ and *Akap11*^-/-^ astrocytes compared to *Akap11*^+/+^. Area of circles represents DEG and DEP counts, defined by FDR < 0.05. (B) Correlation of gene or protein expression in *Akap11*^+/-^ and *Akap11*^-/-^ astrocytes. Pearson’s correlation coefficients (r) are shown on each plot. (C) Correlation of molecular pathway changes between bulk RNA-seq and proteomic datasets in *Akap11*^+/-^ vs *Akap11*^+/+^ and *Akap11*^-/-^ vs *Akap11*^+/+^ astrocytes. Pathways significantly altered at the transcriptomic level (FDR < 0.05) are shown in red, at the proteomic level in blue, and in both datasets in orange. (D) Genes and metabolite products involved in the cholesterol synthesis pathway. (E) Volcano plots highlighting changes in genes involved in cholesterol synthesis pathway (orange dots) in *Akap11*^+/-^ and *Akap11*^-/-^ astrocytes compared to *Akap11*^+/+^. (F) Heatmaps displaying log_2_FC values for all genes/proteins directly involved in cholesterol synthesis. Significance levels are indicated as follows: p-value < 0.05, “°”; padj ≤ 0.1, “.” and padj < 0.05, “*”. **(G-I)** Volcano plots highlighting transcriptomic and proteomic changes in *Akap11*^+/-^ and *Akap11*^-/-^ astrocytes compared to *Akap11*^+/+^, focusing on genes and proteins involved in cholesterol metabolic process (**G**), and fatty acid metabolic process (**H and I**). Orange dots indicate genes within the respective gene sets.

**Figure S2.**
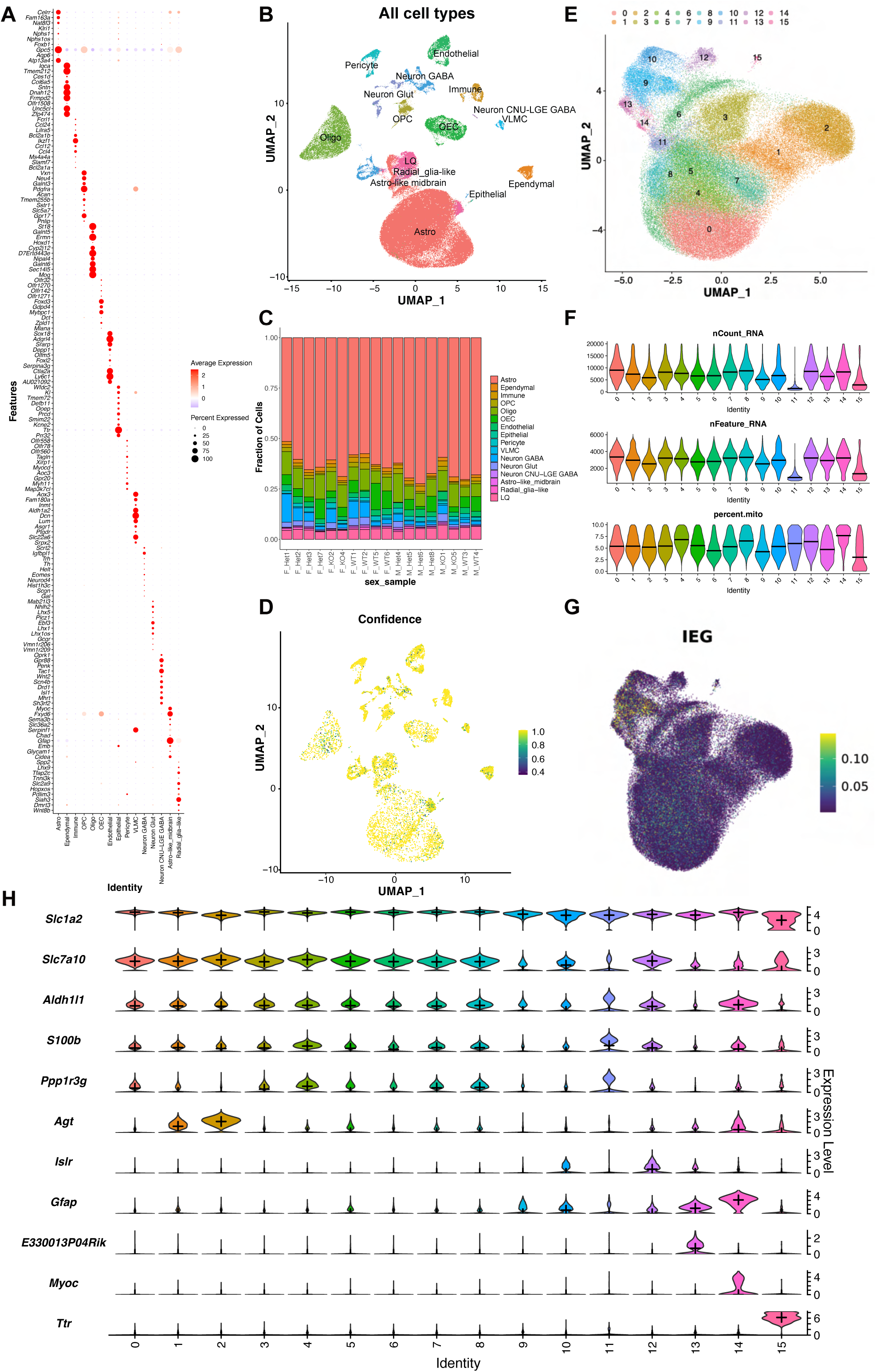
Quality control and annotations of all cell types from scRNA-seq data. **(A)** Dot plot showing the expression of the top 10 DEGs in each cluster (ordered by padj). **(B)** UMAP plot of all cell types identified in scRNA-seq data. **(C)** Bar plot of cell type proportions in each sample. **(D)** Feature plot showing the prediction confidence from automatic cell type labeling (MapMyCells). **(E)** UMAP plot highlighting astrocyte clusters from **(B)**. **(F)** Violin plots displaying the total number of reads (nCount_RNA), the total number of genes (nFeature_RNA), and the percentage of mitochondrial genes (percent.mito) in each cluster. **(G)** Feature plot showing the expression of immediate early genes (IEGs). **(H)** Violin plots showing the expression of astrocyte marker genes.

**Figure S3.**
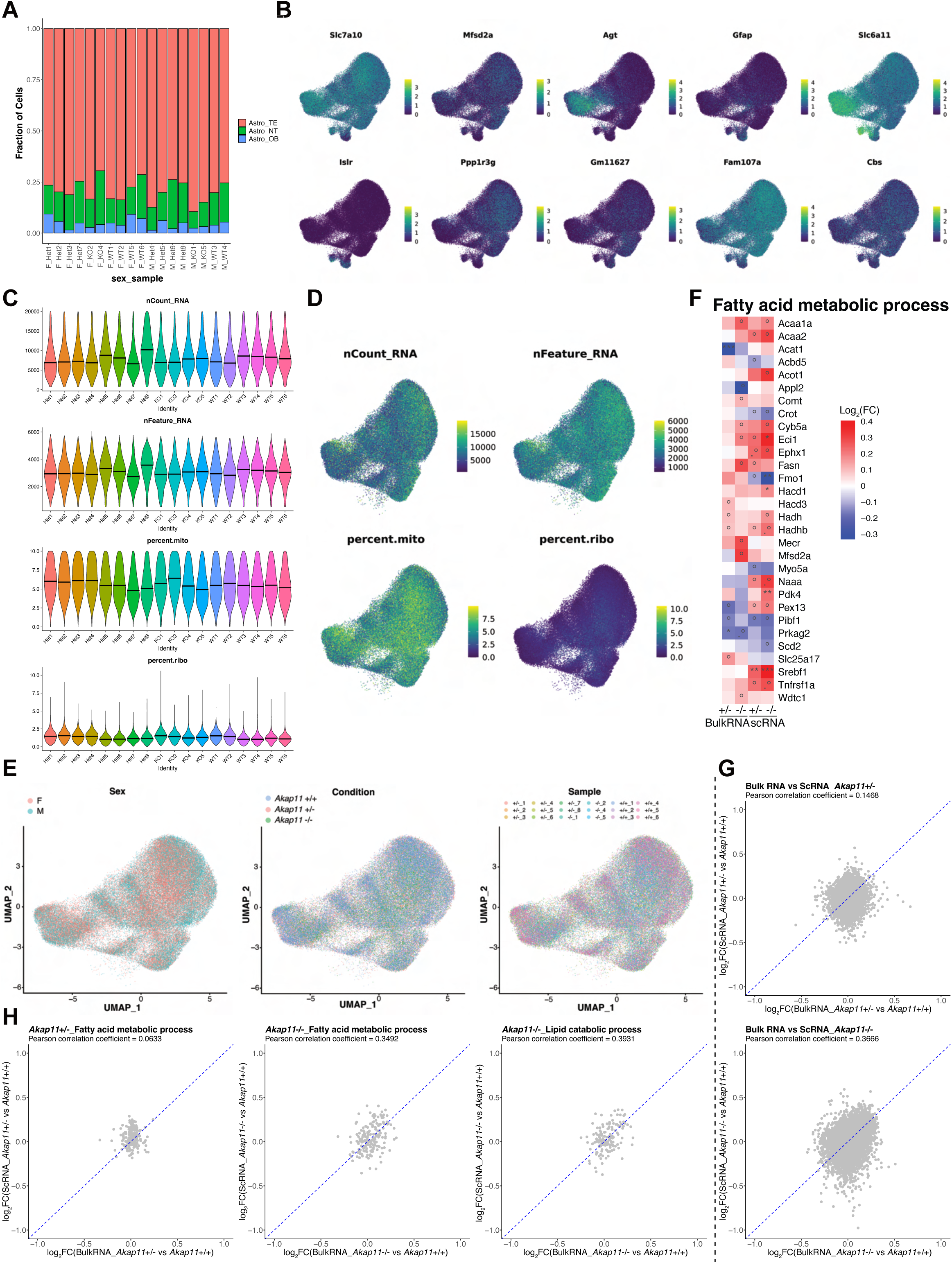
Quality control and annotations of astrocytes from scRNA-seq data. (A) Bar plot showing the proportion of each astrocyte subtype in each sample. (B) Feature plot showing the expression of canonical astrocyte markers. (C) Violin plots displaying the total number of reads (nCount_RNA), the total number of genes (nFeature_RNA), the percentage of mitochondrial genes (percent.mito), and the proportion of ribosomal protein gene expression (percent.ribo) across individual samples. (D) Feature plots showing QC metrics: nCount_RNA, nFeature_RNA, percent.mito, and percent.ribo. (E) UMAP plot of the remaining astrocytes. Cells are color-coded by sex (left), genotype (center), and sample (right). (F) Heatmaps displaying log_2_FC values for genes involved in fatty acid metabolic process. Heatmaps include only those with a p-value < 0.01 in at least one condition. Significance levels are indicated as follows: p-value < 0.05, “°”; padj ≤ 0.1, “.”; padj < 0.05, “*”; padj < 0.01, “**”; padj < 0.001, “***”. (G) Correlation of gene expression between bulk RNA-seq and snRNA-seq data in *Akap11*^+/-^ and *Akap11*^-/-^ astrocytes. Pearson’s correlation coefficients (r) are shown on each plot. (H) Correlation of gene expression for genes involved in fatty acid metabolic process and lipid catabolism process between bulk RNA-seq and snRNA-seq data in *Akap11*^+/-^ and *Akap11*^-/-^ astrocytes. Pearson’s correlation coefficients (r) are shown on each plot.

**Figure S4.**
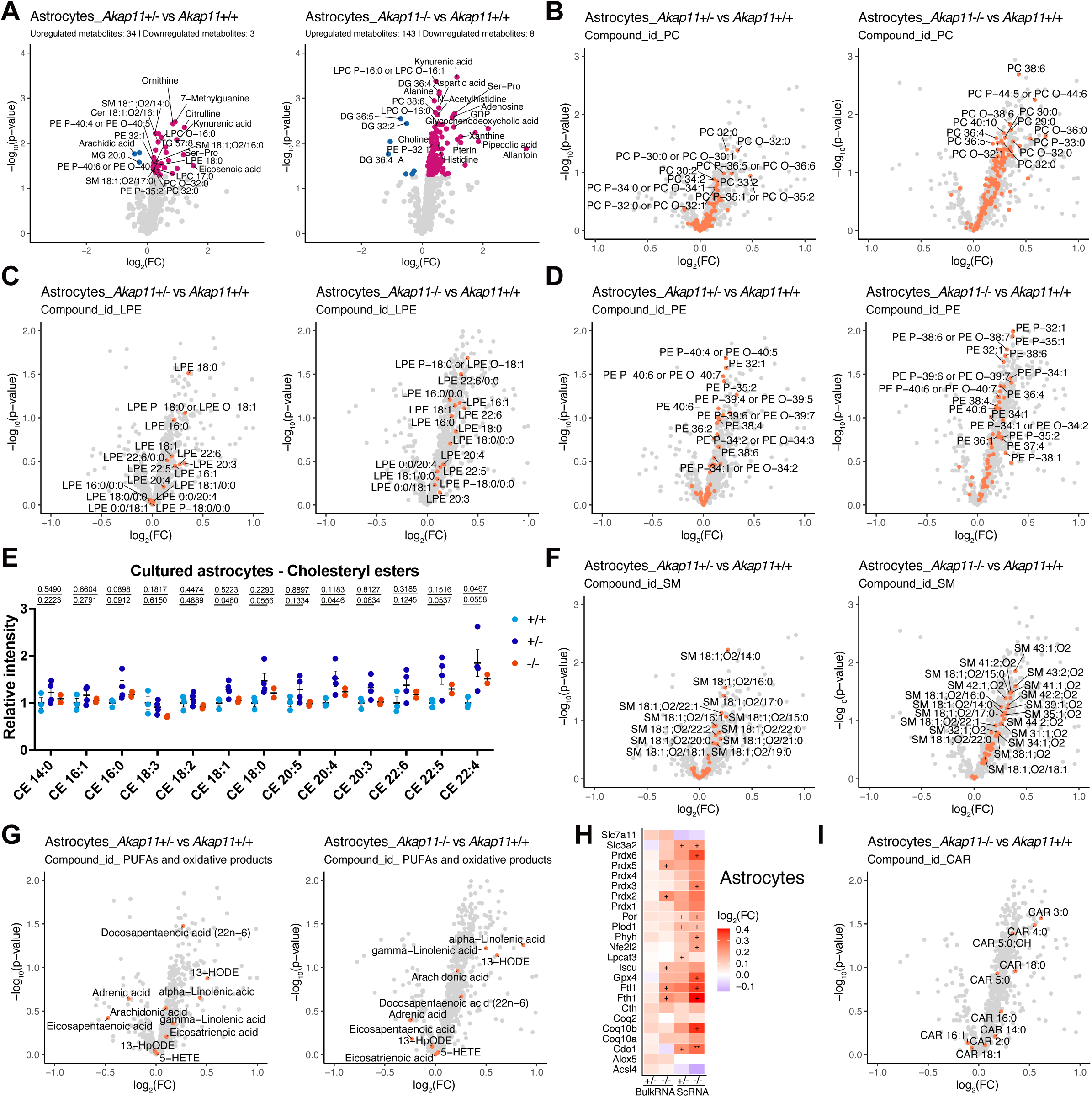
*Akap11* deficiency causes lipids alteration in the cultured astrocytes. (**A)** Volcano plots showing significantly increased (red) or decreased (blue) lipids and metabolites (p < 0.05) in *Akap11*^+/-^ astrocytes (left) and *Akap11*^-/-^ astrocytes (right) astrocytes relative to *Akap11*^+/+^ controls. The horizontal dotted line indicates the significance threshold (p = 0.05). **(B-D)** Volcano plots showing changes in specific lipid species: phosphatidylcholines (PC, **B**), lysophosphatidylethanolamines (LPE, **C**) and phosphatidylethanolamines (PE, **D**) in *Akap11*^+/-^ and *Akap11*^-/-^ astrocytes compared to *Akap11*^+/+^. Orange dots indicate lipids belonging to the respective lipid class. **(E)** Relative abundance of all detected CE species in cultured *Akap11*^+/+^, *Akap11*^+/-^, and *Akap11*^-/-^ astrocytes. **(F)** Volcano plots showing changes in sphingomyelins (SM) in *Akap11*^+/-^ (left) and *Akap11*^-/-^ (right) astrocytes compared to *Akap11*^+/+^ controls. Orange dots represent all identified SM species. **(G and I)** Volcano plots depicting changes in polyunsaturated fatty acids (PUFA) along with their oxidative products (**G**), and acylcarnitines (CAR, **I**) in *Akap11*^+/-^ or *Akap11*^-/-^ astrocytes compared to *Akap11*^+/+^. Orange dots indicate lipids belonging to the respective lipid class. **(H)** Heatmaps displaying log_2_FC values of lipid peroxidation-related genes from bulk RNA-seq of cultured astrocytes and scRNA-seq analysis of mouse astrocytes. Significance levels are indicated as follows: p-value < 0.05, “+”; padj < 0.05, “*”; padj < 0.01, “**”.

**Figure S5.**
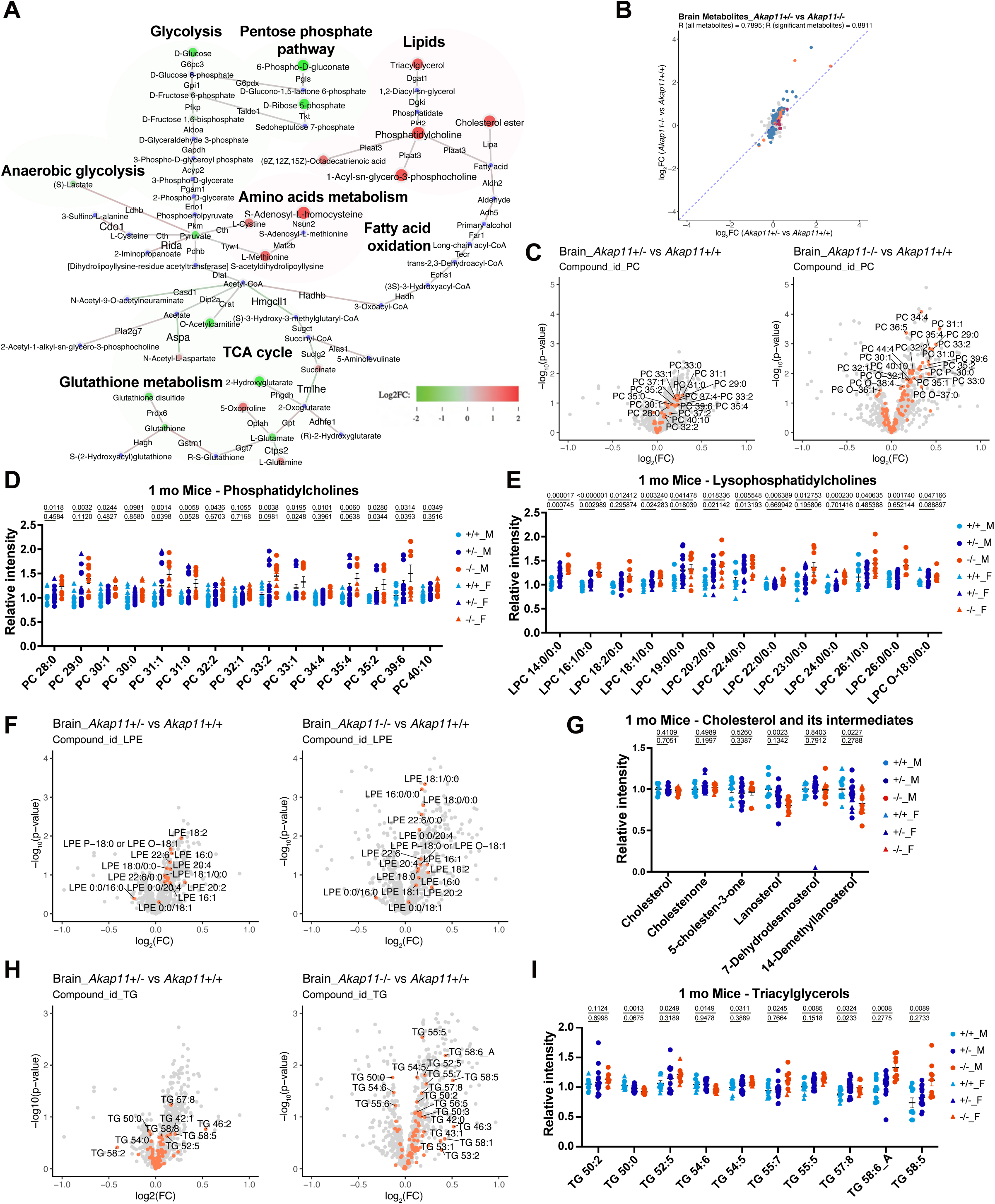
Loss of *Akap11* leads to lipid accumulation in the brain. (**A)** Shiny GATOM integrated analysis of genes and metabolites, combining metabolomic /lipidomic results from brain tissue with scRNA-seq data from mouse astrocytes, highlighting differences between *Akap11*^-/-^ and *Akap11*^+/+^ mice. Nodes represent lipids or metabolites, and edges represent genes. Colors indicate direction of change (red: upregulated, log₂FC > 0; green: downregulated, log₂FC < 0; blue: no significant change). Node size and edge width are proportional to statistical significance (larger = lower p-value). Network visualization was filtered at p < 0.1. **(B)** Correlation of metabolites between *Akap11*^+/-^ vs *Akap11*^+/+^ and *Akap11*^-/-^ vs *Akap11*^+/+^ brain. Metabolites significantly altered (p < 0.05) in the *Akap11*^+/-^ comparison are labeled in red, in the *Akap11*^-/-^ comparison in blue, and in both comparisons in orange. **(C, F, H)** Volcano plots displaying altered lipid species in *Akap11*^+/-^ or *Akap11*^-/-^ mice compared to *Akap11*^+/+^ mice, focusing on PC (**C**), LPE (**F**), and TG (**H**). Orange dots indicate lipids belonging to the respective lipid class. **(D and E)** Relative abundance of representative PC (**D**) and LPC (**E**) species in brain extracts from 1-month-old *Akap11*^+/+^, *Akap11*^+/-^, and *Akap11*^-/-^ mice. (**G)** Relative abundance of cholesterol and its intermediates in brain extracts from 1-month-old *Akap11*^+/+^, *Akap11*^+/-^, and *Akap11*^-/-^ mice. **(I)** Relative abundance of representative TG species in brain extracts from 1-month-old *Akap11*^+/+^, *Akap11*^+/-^, and *Akap11*^-/-^ mice. For **D**, **E**, **G**, **I**, statistical significance was determined by multiple unpaired t test.

**Figure S6.**
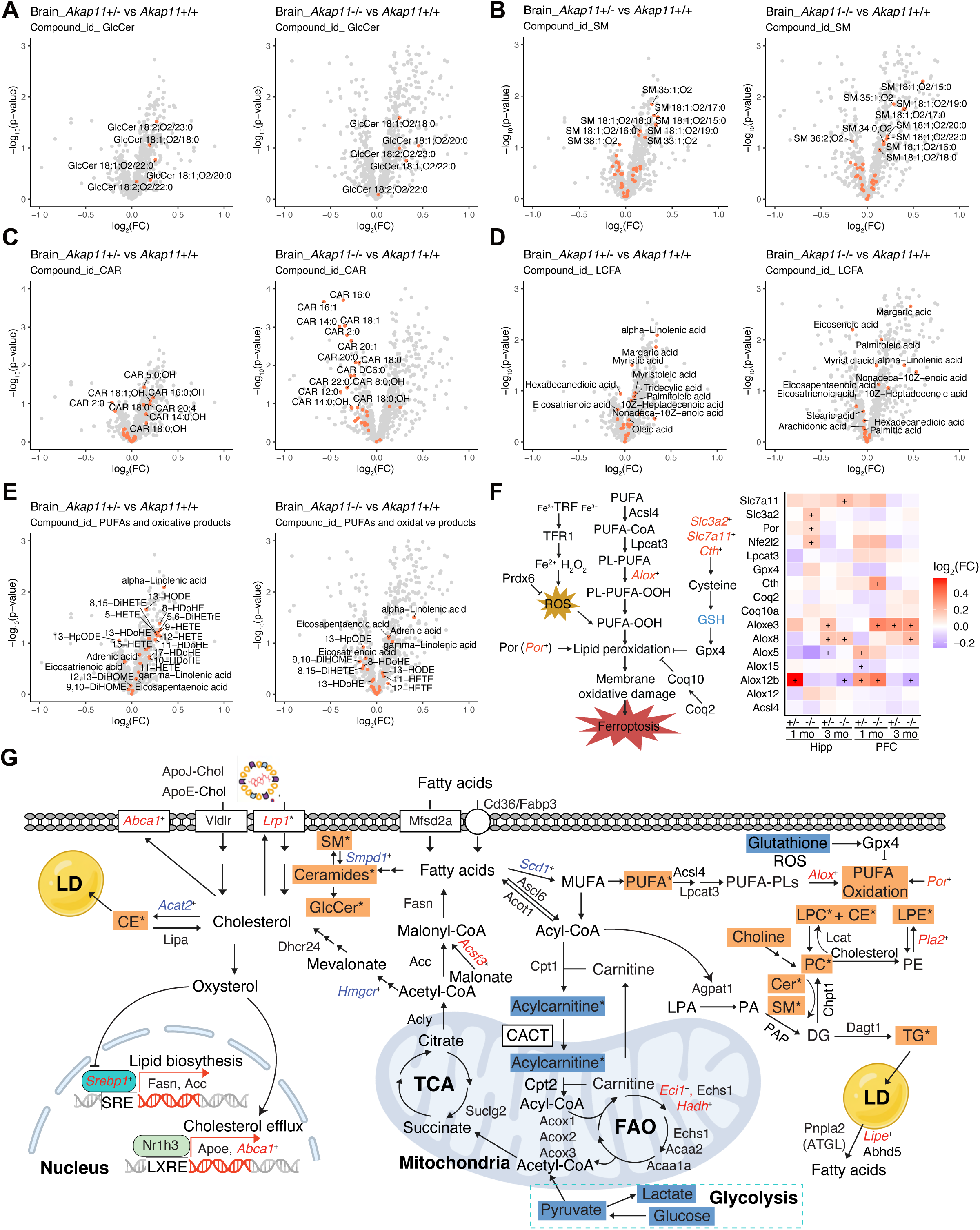
*Akap11* deficiency causes lipids alterations in the brain. (**A-E)** Volcano plots highlighting altered lipid species in *Akap11^+/-^* or *Akap11^-/-^* mice compared to *Akap11*^+/+^ controls, focusing on glucosylceramides (GlcCer, **A**), sphingomyelins (SM, **B**), acylcarnitines (CAR, **C**), long-chain fatty acids (LCFA, **D**), and polyunsaturated fatty acids (PUFA) along with their oxidative products (**E**). Orange dots indicate lipids belonging to the respective lipid class. **(F)** Schematic overview illustrating the lipid peroxidation pathway involved in ferroptosis and corresponding gene expression changes observed in the hippocampus (Hipp) and prefrontal cortex (PFC) of *Akap11* mutant mice. p < 0.05, “+”. **(G)** Integrated analysis of proteomic, metabolomic, and lipidomic data from *Akap11* mutant brain tissues. Differentially expressed genes (italicized) are color-coded (red: increased; blue: decreased) with statistical significance denoted as follows: p-value < 0.05, “+”; padj ≤ 0.1, “.”; padj < 0.05, “*”; padj < 0.01, “**”; padj < 0.001, “***”. Changes in metabolite abundance are indicated by colored rectangles (orange: increased; blue: decreased; p-value < 0.05, “*”).

**Figure S7.**
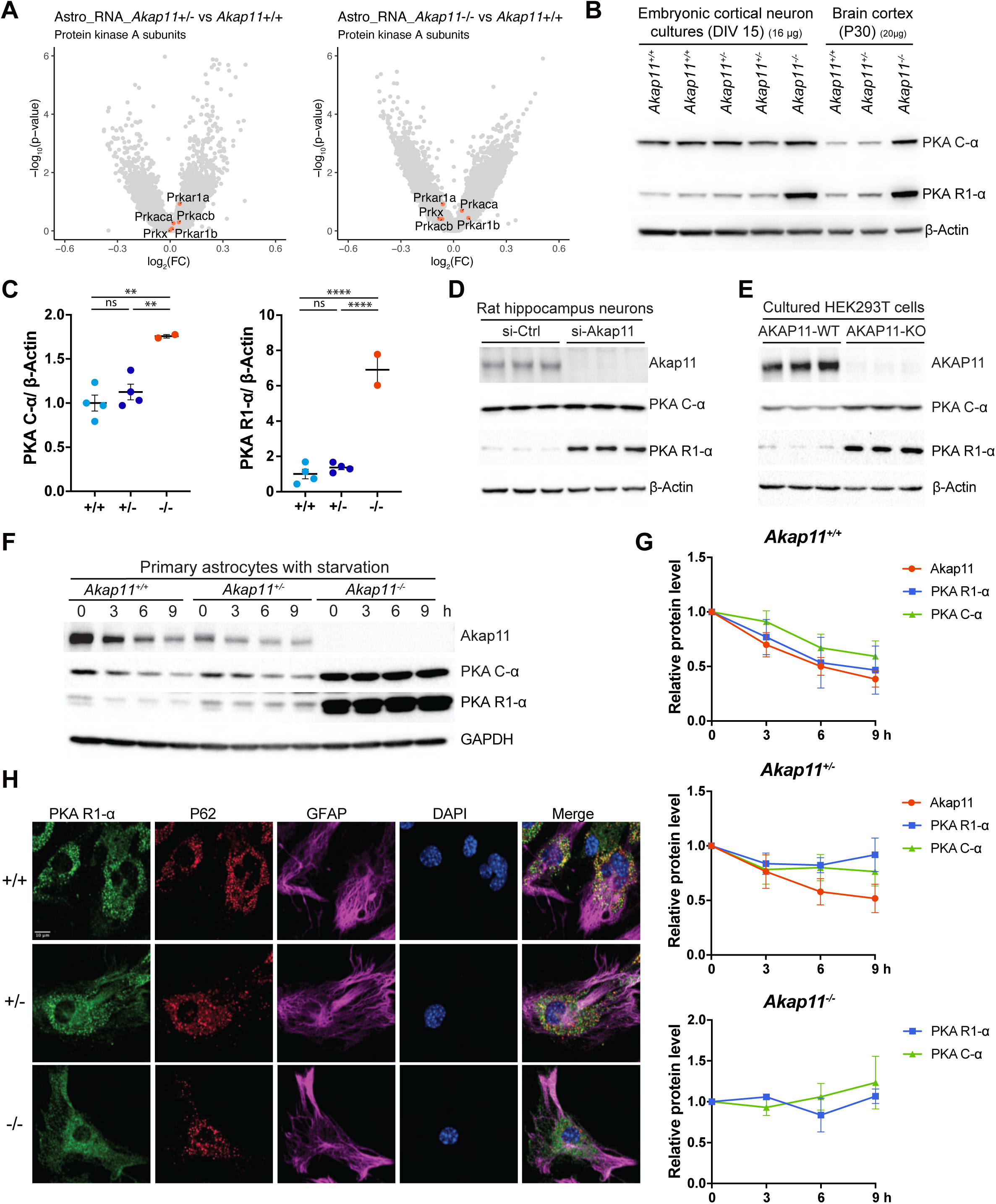
AKAP11 functions as an autophagy receptor for the degradation of PKA subunits. **(A)** RNA levels of protein kinase A subunits remain unchanged in *Akap11*^+/-^ and *Akap11*^-/-^ astrocytes compared to *Akap11*^+/+^ controls. **(B)** Immunoblot analysis of lysates from embryonic cortical neurons and cortical brain tissues of *Akap11*^+/+^, *Akap11*^+/-^, and *Akap11*^-/-^ mice using the indicated antibodies. **(C)** Quantification of the results from (B) with protein levels normalized to β-Actin. Data are presented as mean ± SEM (n ≥ 2 mice/genotype). Statistical significance was determined by one-way ANOVA. *p<0.05; **p<0.01; ***p<0.001; ****p<0.0001; ns, not significant. **(D)** Lysates from control (si-Ctrl) and *Akap11* knock down (si-Akap11) cultured rat hippocampal neurons were analyzed by immunoblotting with the indicated antibodies. **(E)** Lysates from WT and AKAP11 knockout (KO) HEK293T cells were analyzed by immunoblotting with the indicated antibodies. **(F)** *Akap11*^+/+^, *Akap11*^+/-^, and *Akap11*^-/-^ astrocytes were subjected to nutrient starvation in EBSS media for the indicated time points, followed by immunoblotting with the indicated antibodies. **(G)** Quantification of the results from (**F**), with protein levels normalized to β-Actin and then to the time 0 value. Data are from three independent experiments. **(H)** *Akap11*^+/+^, *Akap11*^+/-^, and *Akap11*^-/-^ astrocytes were subjected to nutrient starvation in the presence of Baf A1, followed by immunostaining with anti-PKA R1-α, anti-P62, and anti-GFAP antibodies. Scale bar, 10 μm.

**Figure S8.**
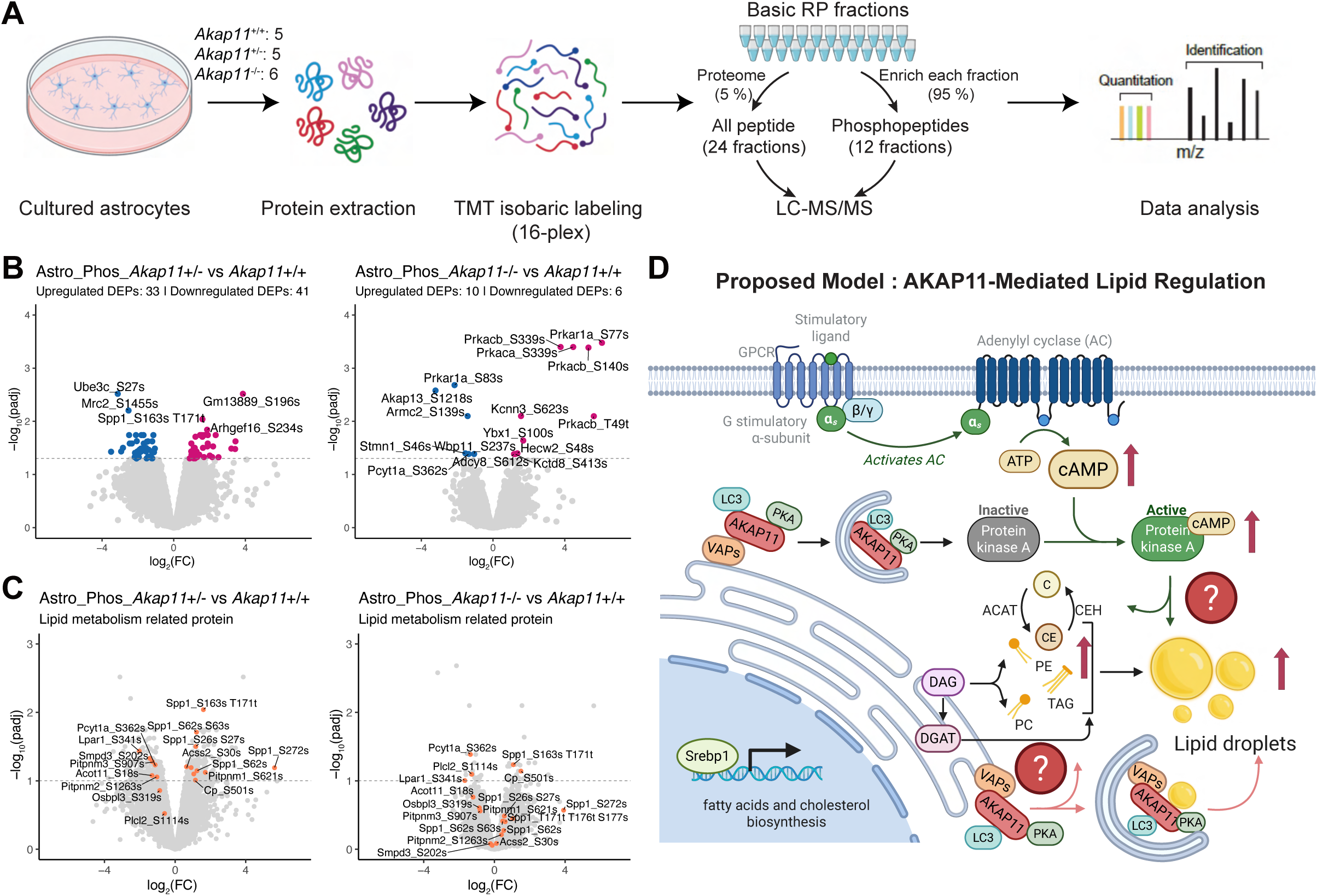
Phosphoproteomic profiling of *Akap11* mutant astrocytes reveals altered phosphorylation of proteins associated with lipid metabolism. **(A)** Schematic overview of the phosphoproteomics workflow. Primary astrocytes were isolated from *Akap11*^+/+^, *Akap11*^+/-^, and *Akap11*^-/-^ mice at postnatal days 1-4 (P1-P4). The same astrocyte preparations were used for the proteomics analysis in Figure 1A. Cell lysates were split, with 5% used for proteomics and 95% for phosphoproteomic analysis. **(B)** Volcano plots showing significantly altered phosphorylation sites (padj < 0.05) in *Akap11*^+/-^ and *Akap11*^-/-^ astrocyte cultures compared to *Akap11*^+/+^ control. The horizontal dotted line indicates the significance threshold (FDR = 0.05). Upregulated and downregulated phosphosites are indicated in red and blue, respectively. **(C)** Selected phosphorylation sites in proteins involved in lipid metabolism were significantly altered (padj < 0.1) in *Akap11*^+/-^ and *Akap11*^-/-^ astrocyte cultures compared to *Akap11*^+/+^ control. **(D)** Proposed model of AKAP11-mediated regulation of lipid metabolism. AKAP11 may modulate lipid homeostasis through the cAMP–PKA signaling pathway. Additionally, its interaction with VAP proteins may influence lipid regulation via autophagy-dependent and - independent mechanisms.

**Figure S9.**
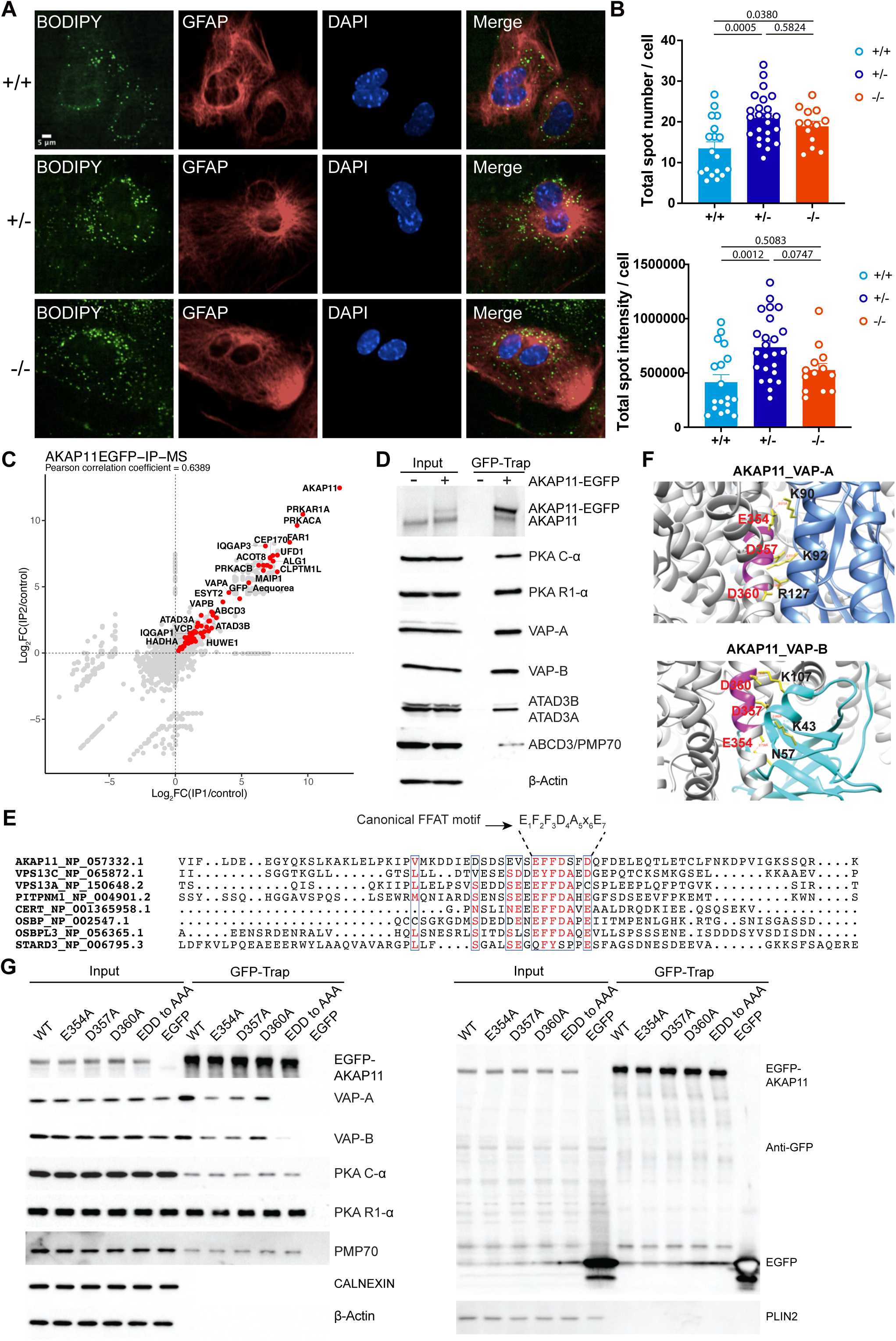
AKAP11 interacts with VAPs through an FFAT motif. **(A)** Cultured astrocytes were stained with BODIPY™ 493/503 (lipid droplets, green) and GFAP (astrocyte marker, red), and DAPI (blue). Scale bar: 5 µm. **(B)** Quantification of the total number of lipid droplets (up) and total spot intensity (bottom) of the BODIPY signal from (**A**). Each dot represents an individual well from cell cultures. The number of mice per genotype used for these experiments: *Akap11*^+/+^, n=2; *Akap11*^+/-^, n=6; *Akap11*^-/-^, n=3. Statistical significance was determined by one-way ANOVA. **(C)** Scatter plot comparing the log₂FC of proteins identified from two independent AKAP11-EGFP immunoprecipitations (IPs) versus control IPs (without AKAP11-EGFP expression) using LC-MS/MS. Proteins significantly enriched in both IPs (p < 0.05) are highlighted in red. **(D)** Validation of selected AKAP11 interactors by immunoblotting. EGFP-tagged AKAP11 was immunoprecipitated using GFP-Trap agarose beads. Total cell lysates (Input) and GFP-Trap precipitates (IP) were analyzed by immunoblotting with the indicated antibodies. **(E)** Amino acid sequence alignment of AKAP11-FFAT and other FFAT-containing proteins using Clustal Omega. **(F)** Predicted interaction interface of AKAP11 with VAP-A and VAP-B, modeled by AlphaFold 3. AKAP11 is shown in grey, VAP-A in blue, and VAP-B in cyan. The AKAP11 FFAT motif is highlighted in magenta. Residues predicted to mediate interaction, including AKAP11 (E354, D357, D360), VAP-A (K90, K92, R127), and VAP-B (N54, K43, R107), are highlighted with yellow side chains. **(G)** Immunoblot analysis of GFP-Trap immunoprecipitates from HEK293T cells stably expressing either EGFP-tagged full-length AKAP11, FFAT motif point mutants, or EGFP control. Blots were probed with indicated antibodies.

**Figure S10.**
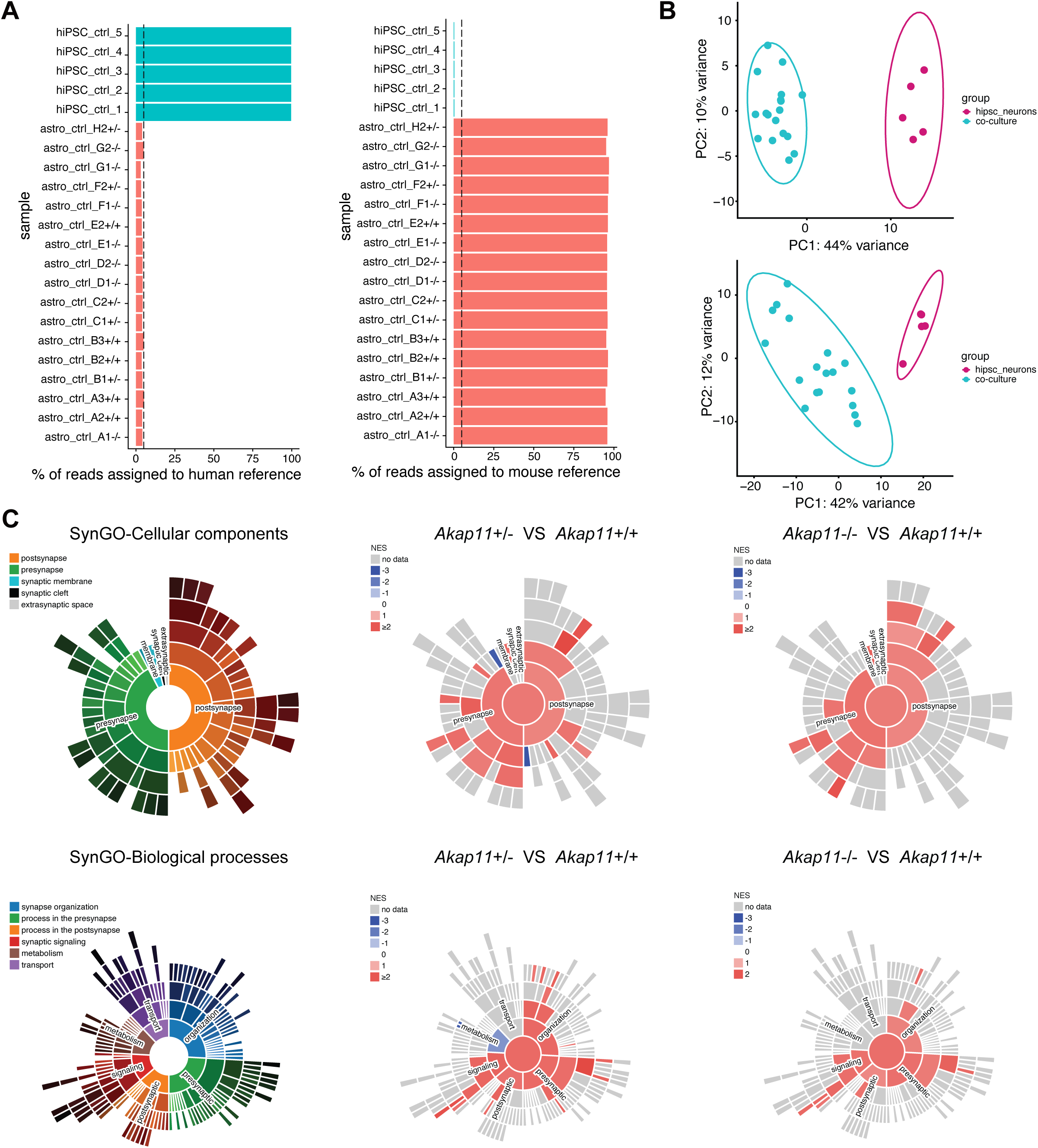
RNA-seq analysis and SynGO enrichment for hiPSC-derived neurons co-cultured with *Akap11* mutant astrocytes. **(A)** Proportion of reads from each sample aligned to the human reference genome and mouse reference genome. **(B)** Principal Component Analysis (PCA) of RNA-seq data from DIV28 neurons (top) and DIV42 neurons (bottom) aligned to a mixed reference genome. **(C)** Sunburst plots showing SynGO terms of cellular components (up) and biological processes (bottom). SynGO enrichment analysis was performed on differentially expressed genes using the SynGO gene set database (v1.1). Significantly enriched synapse-related gene sets (FDR < 0.05) are visualized, with normalized enrichment scores (NES) represented by the color scheme.

**Figure S11.**
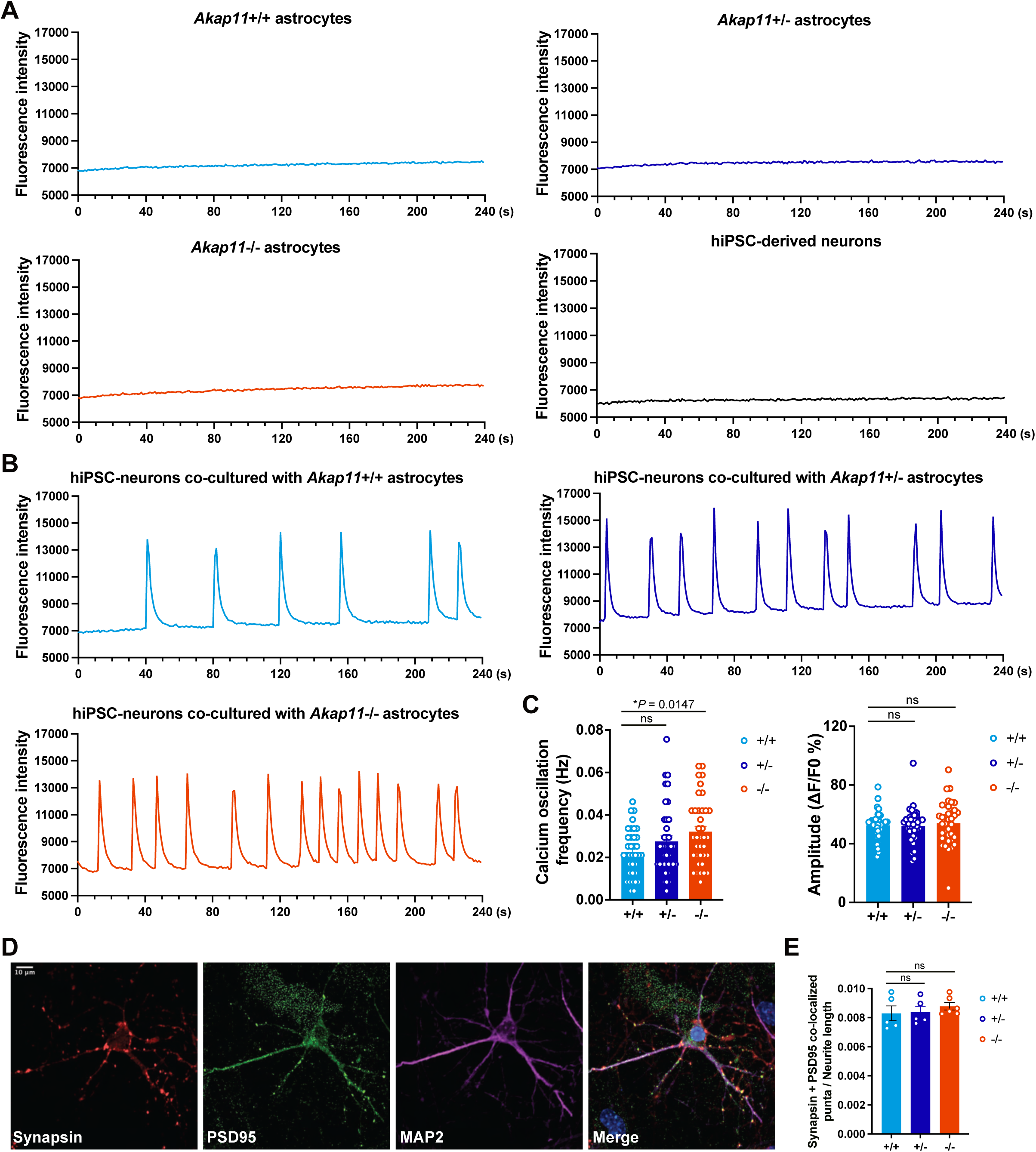
Ca^2+^ oscillations and synaptic density in hiPSC-derived neurons co-cultured with *Akap11* mutant astrocytes. **(A)** Representative traces of Fluo-4 AM fluorescence showing Ca²⁺ activity in hiPSC-derived neuron monocultures and *Akap11*^+/+^, *Akap11*^+/-^, and *Akap11*^-/-^ astrocytes monocultures at DIV42. **(B)** Representative traces of Fluo-4 AM fluorescence showing Ca²⁺ signals in hiPSC-derived neurons co-cultured with *Akap11*^+/+^, *Akap11*^+/-^, and *Akap11*^-/-^ astrocytes at DIV42. **(C)** Quantification of the peak frequency and peak amplitude (ΔF/F₀) of Ca²⁺ oscillations from **(B)**. Each dot represents an individual cell culture. Mice per group: *Akap11*^+/+^ = 5, *Akap11*^+/-^ = 6, *Akap11*^-/-^ = 6. Statistical significance was determined by one-way ANOVA. **(D)** Representative immunostaining of synapses at DIV42 using Synapsin (presynaptic) and PSD95 (postsynaptic) markers. Scale bar: 10 μm. **(E)** Quantification of the density of co-localized Synapsin/PSD95 puncta (number per μm) at DIV42. Each dot represents data from one mouse.

